# Adaptation of mammalian myosin II sequences to body mass

**DOI:** 10.1101/055434

**Authors:** Jake E McGreig, Sarah T Jeanfavre, Charlotte Henson, Michael P Coghlan, Jonathan Walklate, Martin Ridout, Anthony J Baines, Michael A Geeves, Mark N Wass

## Abstract

The speed of muscle contraction is related to body size; muscles in larger species contract at a slower rate. We investigated the evolution of twelve myosin II isoforms to identify any adapted to increasing body mass in mammals. We identified a correlation between body mass and sequence divergence for the motor domain of three adult myosin II isoforms (β, 2A, 2B) suggesting that these isoforms have adapted to increasing body mass. In contrast the non-muscle and developmental isoforms show no correlation of sequence divergence with body mass, while the sarcomeric myosin 7b, extraocular and 2X isoforms showed a divergence intermediate between these two groups. The 2B and β-myosin motor domain showed the greatest rate of sequence divergence (−0.84 and −0.69 % per ten-fold increase in mass respectively). β-myosin is abundant in cardiac ventricle and slow skeletal muscle. We propose that β-myosin has adapted to enable slower heart beating and contraction of slow skeletal muscle as body mass increased.

## Introduction

Striated muscles produce rapid movements of the whole organism (skeletal muscles) or pump blood to transport nutrients and waste products between specialist organs (e.g. cardiac muscle). A key feature of muscle cells is that the speed of contraction (specifically the maximum shortening speed) is a property of the specific myosin isoform(s) expressed in the cell (Pellegrino et al. 2003). The contraction parameters of a muscle adjust to the size of the organism; small animals have fast contracting muscle fibers. As size increases muscle contraction becomes slower to compensate for the greater momentum and inertia associated with a larger mass (Schiaffino and Reggiani 2011) (Hill 1950). This has been widely observed in measured heart rates that are known to be correlated with basal metabolic rate and anti-correlated with body size across a wide range of species (Savage et al. 2007).

The rate of muscle contraction in different species can be controlled in two ways. Firstly, muscles can express different combinations of fast and slow contracting muscle fibres (containing different myosin isoforms) to adjust contraction parameters to physiological requirements (Schiaffino and Reggiani 2011), although most adult muscle fibers express a single myosin isoform (Pellegrino et al. 2003). Secondly, during evolution individual myosin isoforms may adapt to different demand including the change in species size. Here we consider the second of these mechanisms. If the maximum velocity of contraction, driven by a myosin isoform, varies with animal size then there must be specific alterations in the sequence of the myosin to generate the altered velocity.

Myosins are a large family of molecular motors responsible for the generation of forces and movements within eukaryotic cells. Different myosins from the 35 distinct subgroups (Odronitz and Kollmar 2007) organise the actin cytoskeleton, drive cell motility, and participate in organelle & vesicle transport, cell division and signal transduction systems (Krendel and Mooseker 2005). Muscle contraction is driven by members of the myosin II subgroup of the myosin motors. In mammals, the myosin IIs have two branches: one branch includes all of the striated muscle myosins IIs (at least 11 common isoforms) and the second includes both the non-muscle (three isoforms) and smooth-muscle (one gene alternately spliced to give two isoforms) lineage (Golomb et al. 2004) (Schiaffino and Reggiani 2011).

We have investigated evolution of the myosin-II family of proteins in mammals to identify if myosin isoforms expressed in muscle have been adapted to body mass as animals have evolved from smaller to larger sizes. Our data indicate that most fast adult muscle myosin-II isoforms, particularly the β-myosin, have been selectively adapted in relation to mammalian body size.

## Results

All myosin II isoforms contain a head region, or motor-domain, and a tail region/domain. The N-terminal globular motor domain (approximately 800 amino acids) contains all of the requirements for motor activity, while the C-terminal tail domain (~1200 amino acids) drives dimerization and oligomerization into myosin filaments.

Our analysis considered 12 different myosin II isoforms (Table 1), the five main adult sarcomeric isoforms (three fast muscle – 2A, 2B, 2X and two cardiac isoforms, α and β, also known as the slow skeletal isoform), two relatively rare adult sarcomeric forms (extraocular and slow tonic), two developmental isoforms (perinatal and embryonic), a smooth muscle isoform and two non-muscle isoforms (non-muscle a and b). The non-muscle isoforms provide a negative control, as they act at the cellular level and are therefore unlikely to be influenced by body mass. We identified all available complete myosin II sequences of these 12 myosin isoforms (see methods; Table S1). This resulted in a total of 743 sequences, with an average of 61 sequences per isoform (range 41-74 sequences; Table 1). For each isoform about half of the species are from Laurasiatheria (mostly ungulates and cetacean) or Euarchontoglires (rodents and primates) with a couple of Afrotheria or Metatheria. The availability of large numbers of sequences from two separate groups of mammals, Laurasiatheria and Euarchontoglires, allows comparison of evolutionary characteristics after their initial divergence.

**Table 1.** myosin II Isoforms considered

| Gene name | Heavy Chain | Muscle Type | Isoform | Short name | No complete sequences | Motor |  | Tail |  |
| --- | --- | --- | --- | --- | --- | --- | --- | --- | --- |
| Major adult sarcomeric muscle myosins |  |  |  |  |  | R<sup>2</sup> | Gradient (%/log(kg)) | R<sup>2</sup> | Gradient (%/log(kg)) |
| MYH1 | MyHC-1 | Fast | Skeletal 2B | 2B | 41 | 0.838 | -0.84 ± 0.109 | 0.849 | -0.684 ± 0.094 |
| MYH2 | MyHC-2 | Fast | Skeletal 2A | 2A | 74 | 0.442 | -0.544 ± 0.096 | 0.313 | -0.425 ± 0.122 |
| MYH4 | MyHC-4 | Fast | Skeletal 2D/X | 2X | 53 | 0.466 | -0.563 ± 0.205 | 0.52 | -0.71 ± 0.122 |
| MYH6 | MyHC-6 | Cardiac | α cardiac | α | 65 | 0.3 | -0.359 ± 0.285 | 0.324 | -0.279 ± 0.099 |
| MYH7 | MyHC-7 | Cardiac/Slow | β-cardiac | β | 67 | 0.62 | -0.69 ± 0.105 | 0.03 | -0.06 ± 0.164 |
| Specialist adult sarcomeric myosins |  |  |  |  |  |  |  |  |  |
| MYH13 | MyHC-13 | Fast | Extraocular | EXOC | 60 | 0.177 | -0.32 ± 0.101 | 0.214 | -0.717 ± 0.195 |
| MYH7b | MyHC-7b | Slow | Slow Tonic | SlowT | 72 | 0.044 | -0.082 ± 0.059 | 0.016 | -0.051 ± 0.06 |
| Developmental muscle isoforms |  |  |  |  |  |  |  |  |  |
| MYH3 | MyHC-3 | Developmental | Embryonic | EMB | 60 | 0.127 | -0.086 ± 0.035 | 0.224 | -0.313 ± 0.124 |
| MYH8 | MyHC-8 | Developmental | Perinatal | PERI | 49 | 0.009 | -0.039 ± 0.113 | 0.213 | -0.277 ± 0.118 |
| Non sarcomeric myosins |  |  |  |  |  |  |  |  |  |
| MYH9 | MyHC-9 | Non-muscle | Non-muscle A | NMA | 59 | 0.032 | -0.033 ± 0.036 | 0.144 | -0.137 ± 0.061 |
| MYH10 | MyHC-10 | Non-muscle | Non-muscle B | NMB | 66 | 0.111 | 0.042 ± 0.022 | 0.089 | -0.079 ± 0.052 |
| MYH11 | MyHC-11 | Smooth muscle | Smooth Muscle | SM | 71 | 0.498 | -0.356 ± 0.069 | 0.136 | -0.237 ± 0.102 |

MyHC 13 & 7b are labelled *specialist* because unlike the other sarcomeric forms they are not found alone in a specific muscle type but only in combination with other isoforms.

### Adaptation of myosin II motor domain to increasing body mass

To consider how the motor domain of myosin II isoforms may have adapted to increasing body mass, we investigated if there was a correlation between sequence divergence and body mass. Mass values were collected from the literature, and we quote the middle of the range of values reported. The range of masses cover more than six orders of magnitude from 6 g to 10,000 kg (Supplementary Table 1). Being one of the smallest species, mouse was selected as a reference and the sequence identify of the other species with the mouse sequence plotted against the species body mass (see Figure 1 for the β-myosin, the non-muscle 2A and the embryonic isoforms, Supplementary Figure 1 for the other nine isoforms). Sequence identity rather than conservation was used because evolution was considered within a single myosin isoform. A low level of divergence was expected within each isoform and as such even conservative changes of amino acids that tune the function of the protein, may be relevant to adaptation to body mass. For example, a 2-fold change in a rate constant that controls shortening velocity, requires only a change of the order of 1 kcal/mol in the activation energy according to transition state theory. This is comparable to a single hydrogen bond or van der Waals interaction.

**Figure 1.**
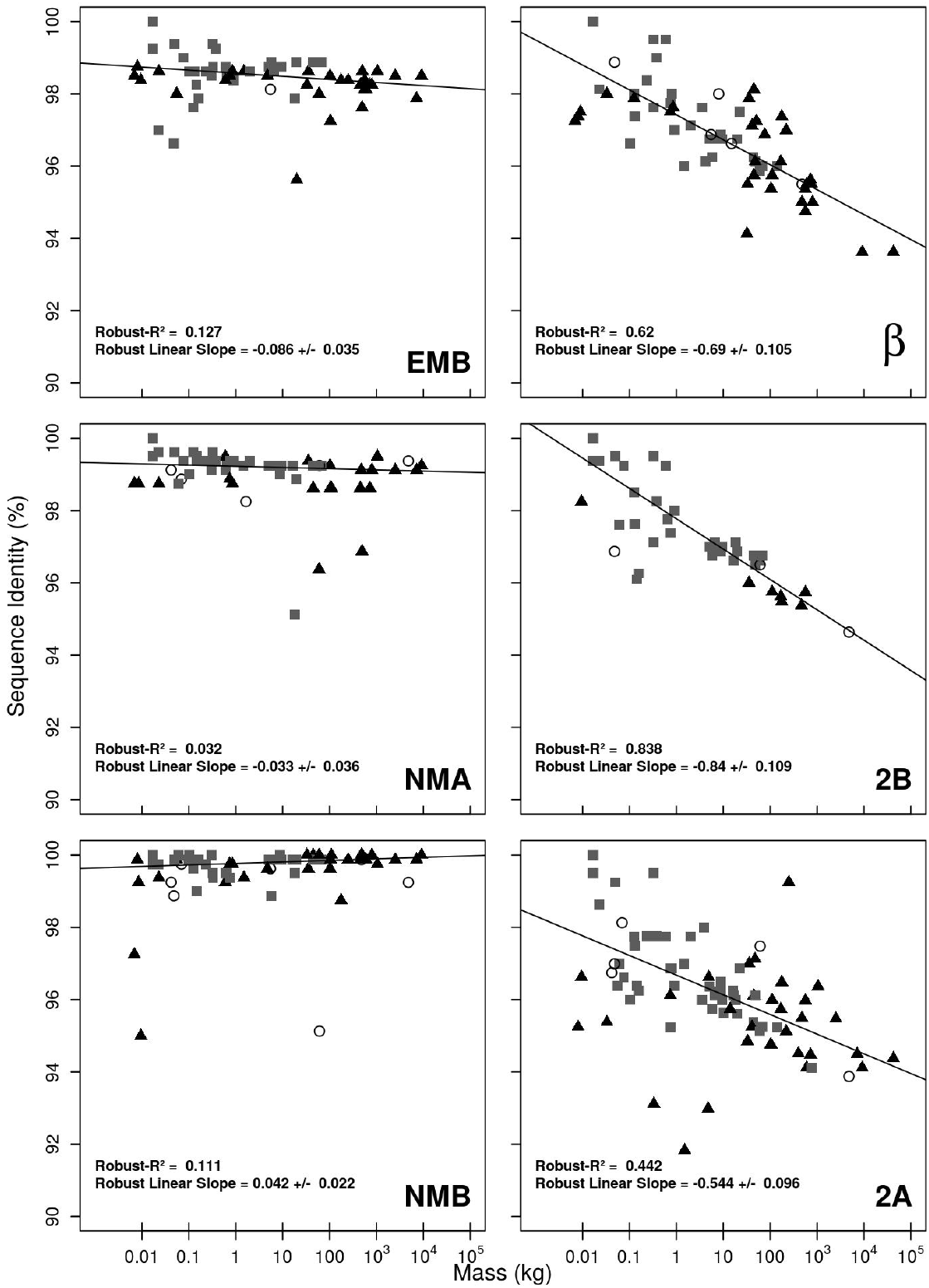
Sequence Identity (%ID) vs Mass (kg) for the motor domains of six myosin II isoforms; embryonic (EMB), non-muscle A (NMA), non-muscle B (NMB), β-myosin, 2A and 2B. Symbols indicate the clade that each species belongs to grey squares (Euarchontoglires), black triangles (Laurasiatheria) and clear circles (Afrotheria and Metatheria). Each plot has been fitted with a robust linear regression. Sequence identity is pairwise to the mouse. The R^2^ value and slope are shown on each plot.

Based on the rate of divergence with body mass, the 12 myosin II isoforms form three groups. The first group contains four of the five main adult sarcomeric isoforms, with the 2B (gradient −0.84 percent divergence per log kg; R^2^ = 0.838, see Figure S1, Table 1) and β-myosin (gradient −0.69 percent divergence per log kg; R^2^ = 0.62, see Fig 2, Table 1) isoforms showing the greatest mass related sequence divergence, followed by the 2A and 2X isoforms (gradients −0.544 and 0.563 respectively; R^2^ 0.442, 0.466), with the 2X isoform plots showing a wider scatter in data as indicated in the 20-40% error in the gradient value.

**Figure 2.**
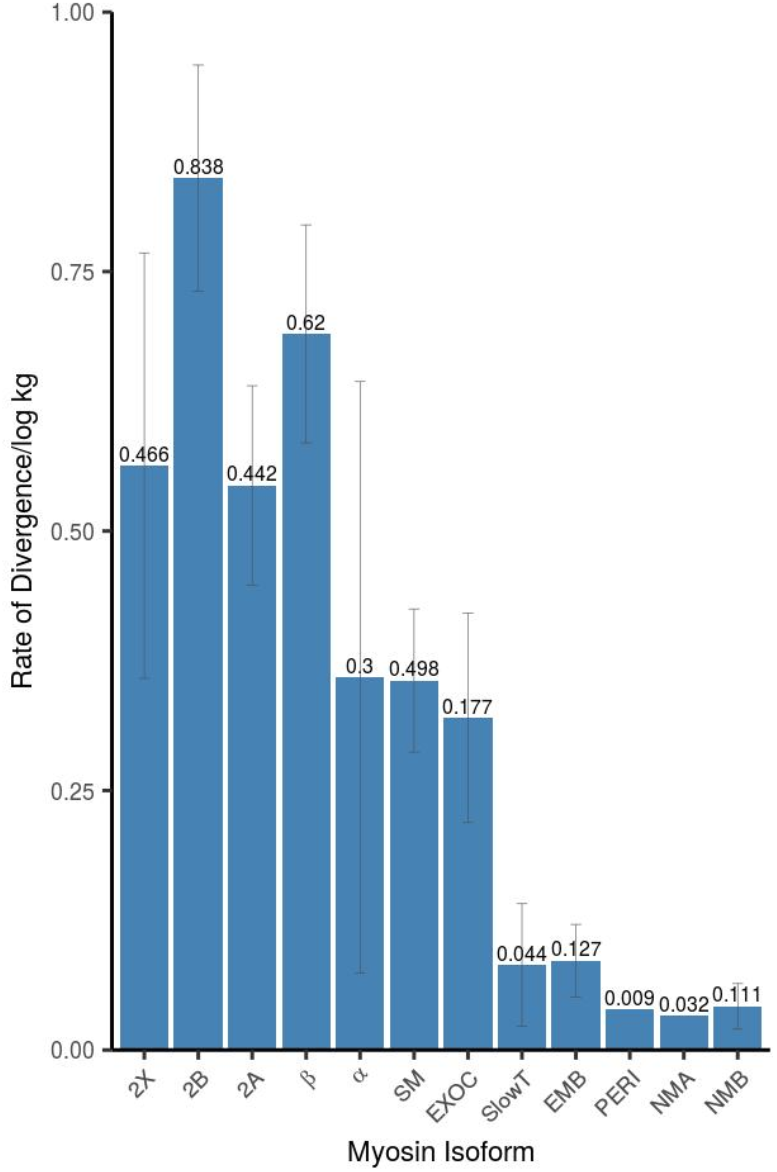
The relationship between body mass and sequence divergence for the Myosin II motor domains. The rate of sequence divergence per log kg is plotted for the 12 myosin II isoforms. Values above the bar are the R^2^ values. The error bars show the error of the gradient.

At the other extreme five of the isoforms (slow tonic, embryonic, perinatal, non-muscle A and non-muscle B) exhibit little divergence of sequence with body mass (slope −0.086 – +0.042; highest R^2^ = 0.13).

This result with quite distinct dependence upon mass suggests different selection conditions apply to the two groups. This is compatible with the adult muscle myosins adapting to the requirements of changes in size whilst the cellular myosins were not exposed to this selection pressure and were highly constrained in the sequence.

The remaining three isoforms (α-myosin, smooth muscle and extraocular) are intermediate between the two groups with a lower rate of sequence divergence with body mass (−0.3 – −0.4) with only the smooth muscle isoform having an R^2^ > 0.4 (0.498).

### Adaption of β-myosin motor domain to reduce heart rate as species size increased

We focussed on β-myosin as it is primarily expressed in cardiac muscle, and therefore performs the same specific function in different species. Whereas other striated muscle isoforms are expressed in multiple tissues and may be involved in multiple different functions. The negative correlation between species heart rate and body mass is well established (Savage et al. 2007). Given our observation that β-myosin shows the greatest association of sequence change with body mass, it is possible that there has been selection pressure on this isoform to enable the change in heart rate as body mass has increased.

To investigate this hypothesis we selected the β-myosin and two other isoforms, the embryonic and non-muscle A (NMA) as controls, where we do not see a relationship between sequence divergence and body mass (Figures 1,2, Figure S1). These three isoforms were considered using a phylogenetic framework. Phylogenetic trees (Figure 3 and Supplementary Figure 2) were generated using both Bayesian and maximum likelihood approaches (see methods). Both methods produced trees that were essentially consistent with each other, with the tree comparison scores for embryonic, β-myosin, and non-muscle A (NMA) isoforms being similar at 0.78, 0.83, and 0.84 respectively. The β-myosin reflects known evolutionary patterns (Figure 3) with the exception of *Trichechus manatus latirostris*, from the Afrotheria clade, which is grouped with the Euarchontoglires. The trees for both the embryonic and NMA isoforms do not clearly separate the two main clades, which may not be surprising given that most of the sequences are more than 99% identical (Figure 1). Nonetheless, these overall data provide a strong phylogenetic framework for further analyses.

**Figure 3.**
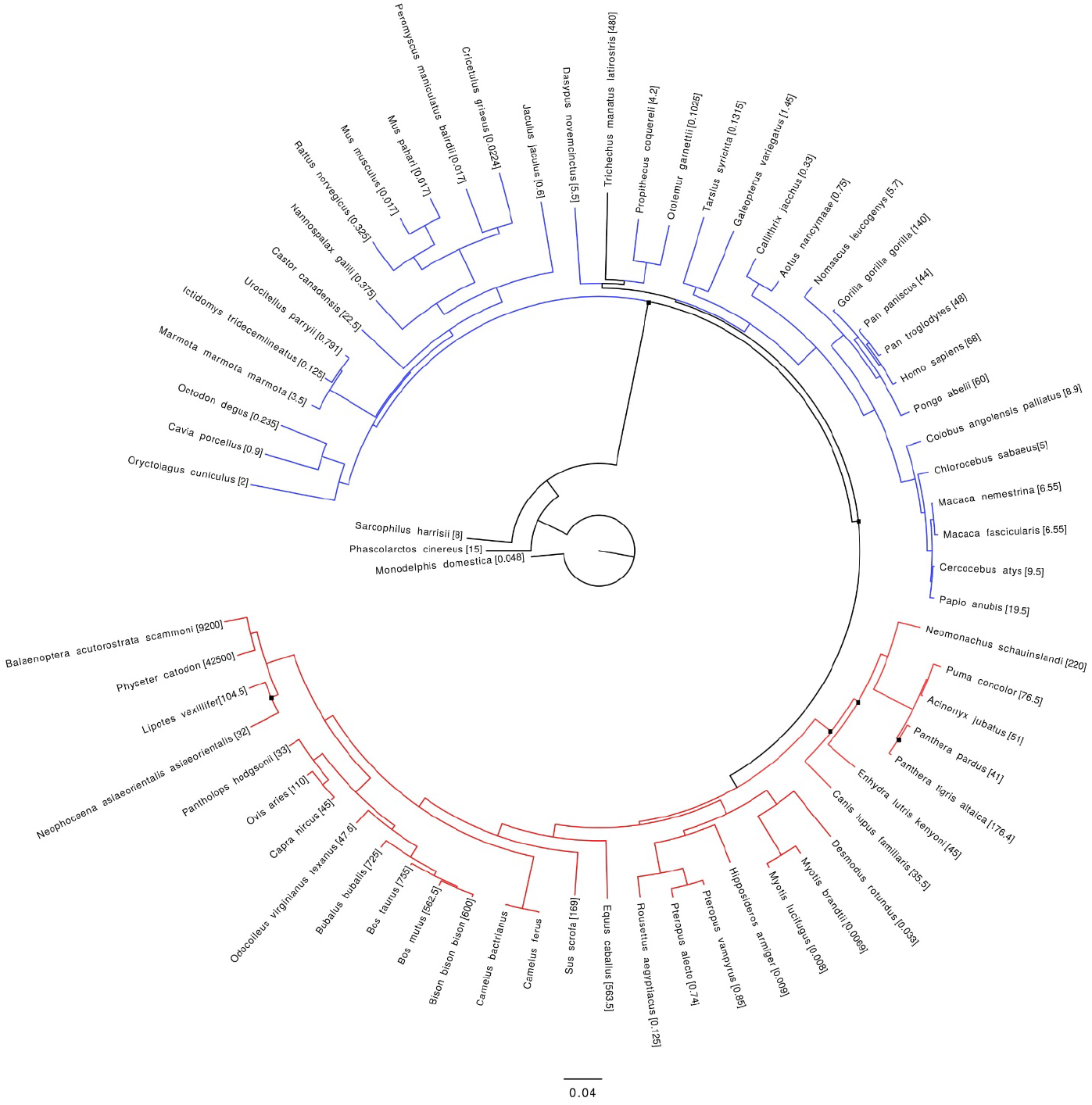
Phylogenetic Tree for the coding DNA sequences of the β-myosin (MyHC 7) motor domain. The Maximum likelihood tree is shown. Black circles indicate positions where the Bayesian tree differed. The tree is coloured according to clade: Blue lines are Euarchontoglires (blue lines), Laurasiatheria (orange) and the few species in other groups are coloured black. The species mass is shown in brackets with the corresponding species name; all have the unit kg.

To consider if the sequence divergence observed in β-myosin is a result of selective pressure, we examined the rate of change at the DNA sequence level. We calculated pairwise dN/dS ratios (Non-synonymous rate/synonymous rate; each species compared to mouse) for the three myosin isoforms (Figure 4 and Supplementary Figure 4). For the β-myosin motor domain there is a clear correlated increase in dN/dS with increasing body mass (Figure 4). This correlation is due to increasing non-synonymous changes, while the rate of synonymous changes remains relatively stable (Figure 4). As the rate of synonymous changes was constant and independent of body size, this implies that there has not been a general increase in the rate of mutations in the nucleotide sequence, instead there has been a specific increase in non-synonymous changes. Moreover, the patristic distances for all the β-myosin motor domains, reflecting total nucleotide changes, show no correlation with body mass (Supplementary Figure 3). A similar dN/dS ratio was observed for the fast muscle isoforms 2A, 2B and 2X (Fig S5). Together, these results demonstrate a selective pressure on β-myosin and the three fast muscle myosins to accept protein sequence changes and these appear to enable a change in contraction speed (and heart rate for β myosin), as body mass changes. In contrast, neither the embryonic nor NMA isoforms showed any correlation in dN/dS with increasing body mass, where the ratio was relatively constant (Figure 4).

**Figure 4.**
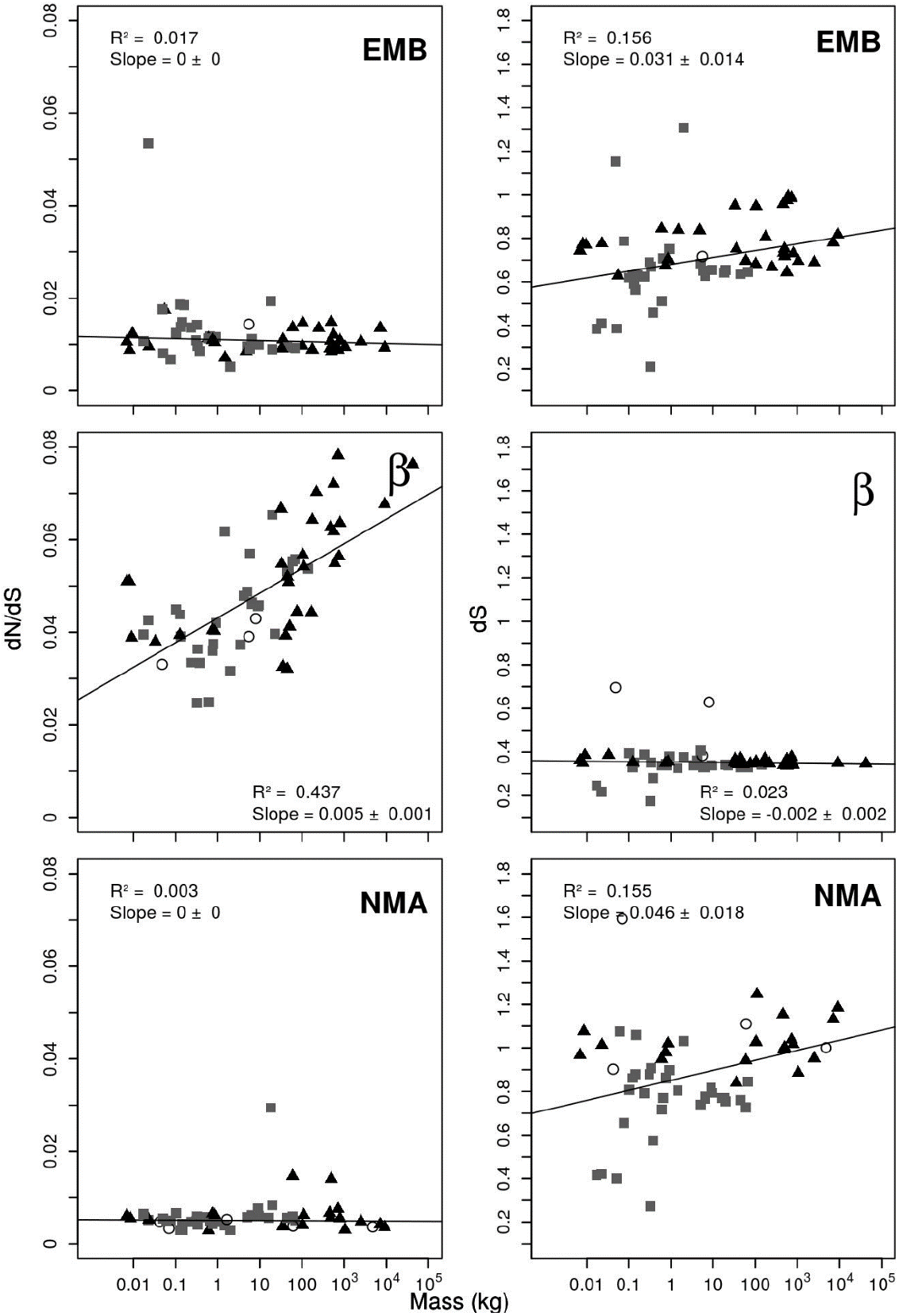
DNA sequence indicators of positive selection on species mass for three myosin heavy chains. Right panels: % synonymous (dS) sequence identity. Left panels The Non-synonymous (dN) to synonymous (dS) variant ratio (dN/dS) to the mouse sequence for embryonic, beta, and non-muscle A myosin as a function of species mass. Details of the species mass and sequences used are given in Table S1. Markers indicates clade: Euarchontoglires (grey squares), Laurasiatheria (black triangles) and Afrotheria and Metatheria (open circles).

### Comparison of myosin species variation with human variation in gnomAD

To further investigate if the divergence observed in β-myosin is related to changing species heart rate we considered the positions in β-myosin that vary between species (i.e. an alternative amino acid present in at least six species) with both known variation in human β-myosin from the gnomAD study (Lek et al. 2016; Karczewski et al. 2019) (variation data for a population of over 140,000 individuals) and positions at which mutations associated with human cardiomyopathy occur (Figure 5).

**Figure 5.**
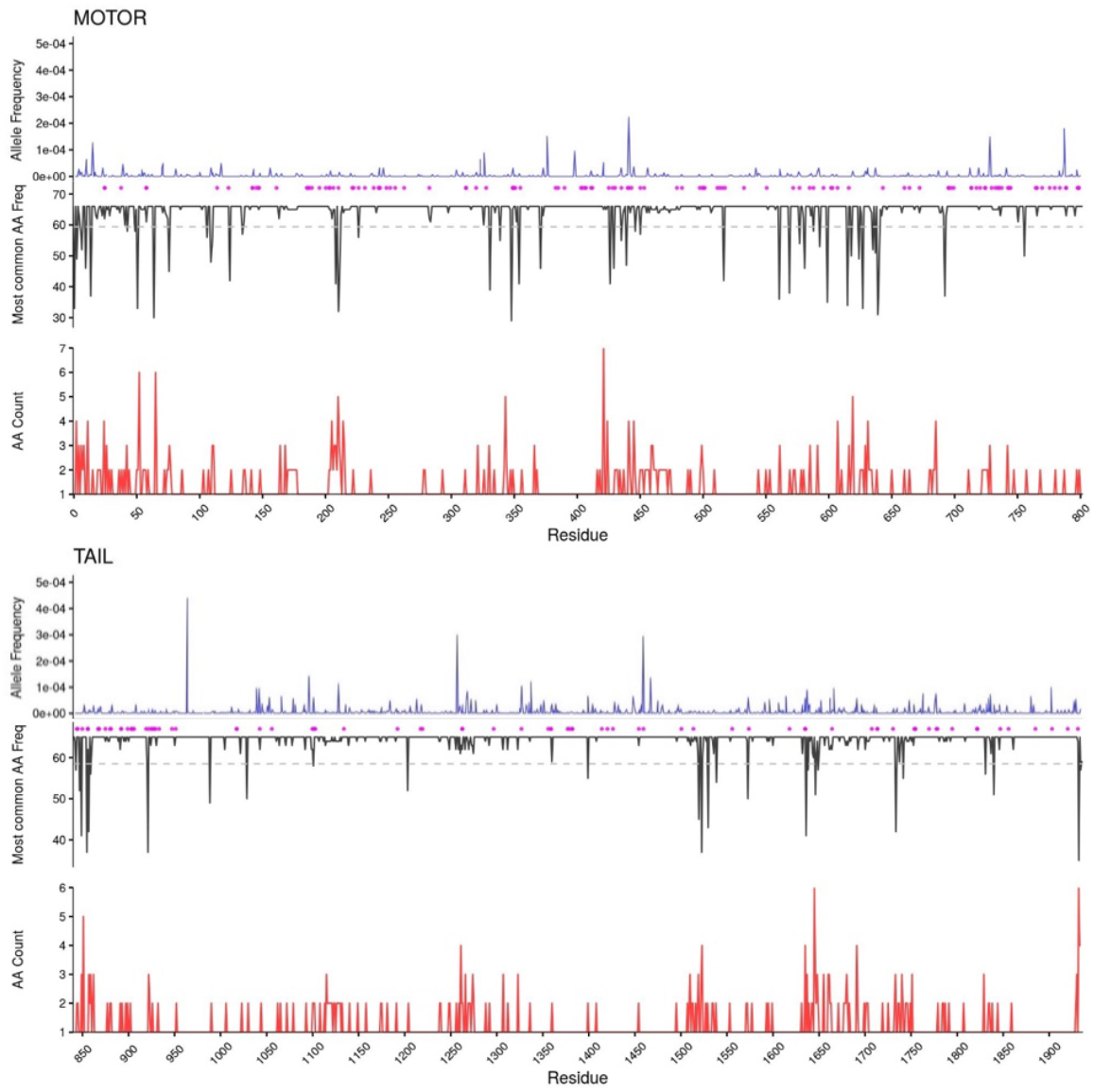
Location of human variation, cardiomyopathy associated variants and nonconserved residues for the β isoform._Blue lines show the frequency of missense variation present in the Genome Aggregation Database (GnomAD). Magenta circles indicate the residue position of variants associated with cardiomyopathy (extracted from UniProt). Black lines show the number of times the consensus amino acid occurs at each residue position in the sequence (maximum equals the no of sequences used). Red lines shows the number of different amino acids occurring at that position in the sequence (minimum 1). The dashed grey lines indicate the position at which more than 10% of sequences contain an alternate amino acid.

There is very little variation in the motor domains of the β-myosin, embryonic and NMA isoforms within the human gnomAD population (Figure 5A, Supplementary Figure 6), with allele frequencies ranging from 0 to ~2×10^-4^. Variation is observed at a higher percentage of positions in the β-myosin tail, 50% of residues compared to 30% in the β-myosin motor domain. Also there are more higher frequency variants in the tail than the motor (Figure 5). These observations suggest that variation in these motor domains is largely not tolerated within humans. In our analysis comparing different species we also see that the motor domains of the embryonic and non-muscle A isoforms are highly conserved (20 and 10 positions that differ in more than 10% of sequences respectively – Figure S6). So the motor domains of these isoforms are highly conserved both within human and between species. In contrast while the β-myosin motor domain is highly conserved within humans, there is much greater variation between species for motor domain with 50 positions that differ in >10% of sequences (Figure 5).

Mutations in β-myosin are known to cause cardiomyopathies, with 103 residues in a domain of 800 associated with the disease (Buvoli et al. 2008; Walsh et al. 2010; Moore et al. 2012). This further suggests that the β-myosin motor domain sequence in humans is therefore highly constrained. In contrast, our analysis shows that there is much greater variation between species (Figure 5A – red line). Four of the 53 positions that vary in >10% of species coincide with known cardiomyopathy positions (residues 211, 222, 430, 797) and a further 12 are within three residues of cardiomyopathy positions (residues 208, 210, 214, 326, 424, 434, 569, 573, 585, 607, 616) demonstrating that some changes that cause disease in humans have been selected for in other species. Together with the gnomAD data this suggests that the variation observed between species is likely to be functionally relevant, as the sequence is highly conserved in individual species, as demonstrated for humans where many positions of variation are associated with disease.

### Comparison of the β-myosin motor and tail domains

The motor domain, as the name suggests, is the part of the myosin responsible for myosin motor activity and has therefore been the focus of the analysis; this domain determines the speed of contraction. The analysis of sequence divergence as a function of body mass was also considered for the tail domains of all of the myosin isoforms. This provides a further control in addition to the analysis of the motor domains of the embryonic and NMA isoforms. The β-myosin tail domain sequence had a much lower divergence with body mass than the motor domain (gradient −0.06 ±0.164, R^2^ 0.03; Figure S1, Table 1) and the dN/dS ratio did not change with increasing body mass (Figure S5). With the exception of the two of the fast muscle myosin isoforms 2B & 2X the R^2^ values for the tail domain were < 0.4 for all isoforms, indicating little correlation of sequence change with mass. For the fast muscle isoforms (2B, 2X) tails the R^2^ values and the slopes indicate a similar dependence on mass as for the motor domain (Figure 1, S1). This correlation was also seen in the dN/dS data (Figure S3, S4).

### Location of sequence changes in β-myosin

The location of sequence changes in the motor and tail domains were examined to evaluate if particular structural features of the domains are especially variable. The location of each of the sequence changes observed for the β, non-muscle A and embryonic myosins were analysed (Figure 5,S6). For each residue we considered the frequency of the consensus amino acid (black lines) and the number of different amino acids present at each position (red lines). In the β-myosin motor domain, of the 800 amino acids 632 were totally conserved and a further 114 sites were highly conserved (i.e. fewer than four species have a different amino acid, for 69 of these sites only one species had a different amino acid). These changes occurred in so few species that no conclusions could be drawn about the driver for these changes. For fifty-three positions the consensus amino acid was present in less than 90% of the sequences, thus these positions with more frequent alternate amino acids are of greater interest (highlighted in Figure 5 by crossing the dotted line).

The sequence variations are scattered throughout the motor both within and outside the major functional regions of the motor domain with no identifiable pattern to the location of the changes (Figure 5). While 15% of the positions that vary between species correspond to human myopathy positions, there is no simple correlation between the two groups of positions. For example, cardiomyopathy linked mutations are enriched in the 68 amino acid converter region (Homburger et al. 2016)(residues 710-778) but there are very few species sequence changes in this region with only one position that differs in more than two species (Gly747, replaced by Ser in seven species).

Complete analysis of the sequence changes and their possible functional significance is beyond the scope of the current work. A detailed bioinformatics and experimental analysis of the functional consequences of the sequence changes in β-cardiac myosin will be published separately (Johnson et al. 2019). The type of issues raised by this analysis can be illustrated by consideration of Loop1. This is a surface loop near the entrance to the nucleotide-binding site, that is hypervariable in both length and sequence between myosin classes and has been associated with controlling ADP release from myosin (Spudich 1994; Goodson et al. 1999). The rate constant for ADP release has been implicated in setting the maximum shortening velocity in some myosin IIs (Bloemink and Geeves 2011). In some species and muscle types the myosin gene is alternatively spliced in this region to generate myosin with different functions e.g. vertebrate smooth and scallop muscle myosins (Kurzawa-Goertz et al. 1998; Sweeney et al. 1998).

In the β-myosins analysed here, Loop 1 is identical in length throughout, with the consensus sequence ^202^GDRSKK<u>D</u>Q<u>TP</u>GKG^214^ and is highly conserved. Only the three underlined sites have alternate residues (Figure 6). While some patterns are observed, there is no simple relationship between residues changes and species size for this region. For example, residue 208 changes between Asp and Glu with Glu predominant in larger species and Asp in smaller. Multiple amino acids are observed at residue 210 but Asn and Thr predominate, with Asn primarily in the smaller species (not present in any of the 30 largest species). Pro-211 is replaced in 23 species by Thr or by Ser, yet in the 25 smallest species Pro is predominant (occurs 23 times). A switch of amino acid between small and large species is not common to all of the 53 sites with alternative amino acids but such a pattern can be discerned in several locations for both conservative and non-conservative substitutions.

**Figure 6.**
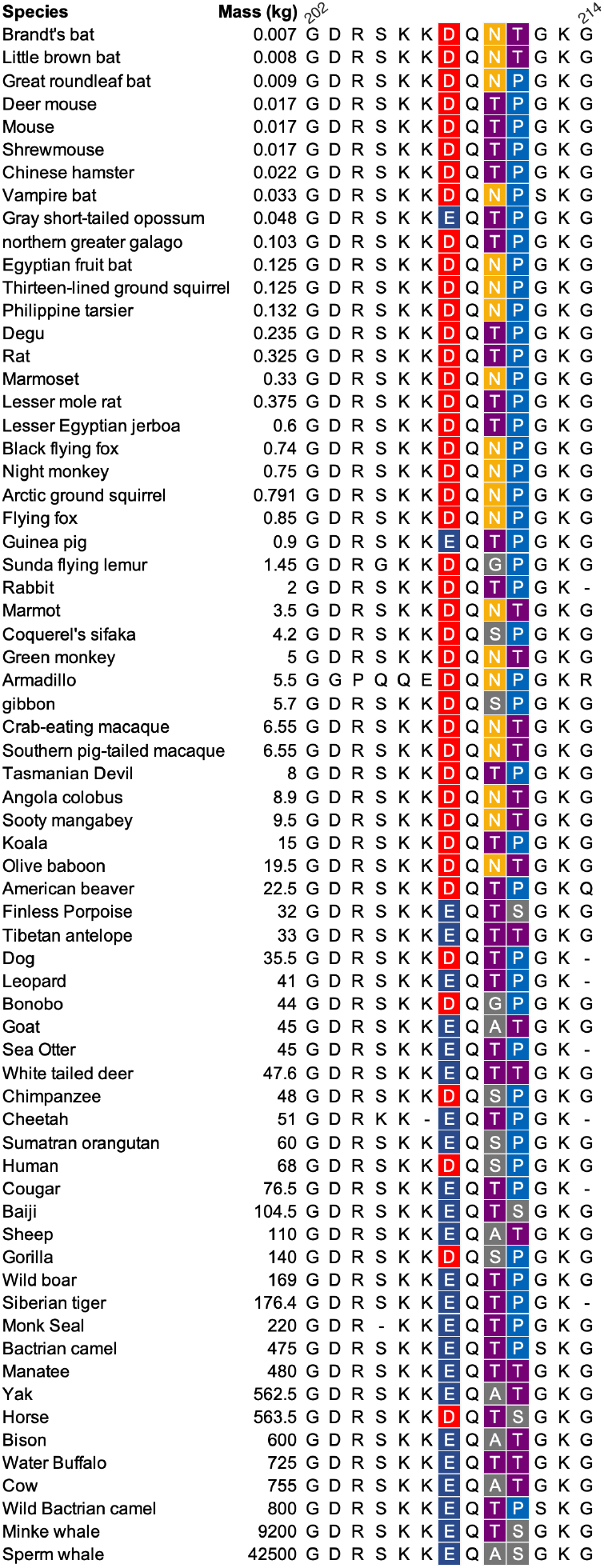
Variation in the β-myosin loop 1 region. Loop one is shown for the 67 β-myosin sequences from smallest to greatest body mass (kg). The loop contains three positions (208, 210 and 211) where there is considerable variation and there is a trend for one amino acid in smaller species and a different one in larger species.

In the other two myosin isoforms there are very few changes in sequence in the motor domains (Supplementary Figure 6). Very few of these have more than a single alternate amino acid. The sequence changes in the tail are much more common and are too frequent to pick out any specific details (Supplementary Figure 6).

## Discussion

Our analysis of myosin evolution has identified that non-muscle and developmental myosin II isoforms have not adapted to increased species size, while there is strong evidence for adaptation in the β-myosin and two adult fast myosin II isoforms (2A, 2B) motor domains whose sequence divergence is correlated with body mass. The fast muscle 2X and Smooth muscle and α isoforms are intermediated.

The myosin motor domain directly controls muscle contraction velocity and we have shown that adaptation to changes in body mass occurred via variation in the myosin motor domain of several adult muscle isoforms (β, 2A & 2B). We propose that the variation identified in the β-myosin motor domain is required to optimize the myosin for the required contraction velocity/heart rate. In the β motor domain we observed a 0.69% change in sequence per 10-fold change in mass (kg). For a motor domain of 800 amino acids this translates to approximately 5.5 amino acids for each 10-fold change in mammal mass.

Our analysis suggests that two of the other main human myosin II isoforms (2A, 2B) have also adapted to increasing body mass. A complexity of studying mammals is that they do not all use their muscles and myosin isoforms in the same way or for the same purpose. Gait and posture, for example, vary considerably and may contribute different selective pressures to muscle and myosin divergence. Mammals are remarkably adaptable and muscle tissue is particularly plastic. Additionally, the contraction parameters (including velocity) of a skeletal muscle can be adapted by adjusting the mix of fiber types (and hence myosin isoforms) used in a specific muscle. Thus, the selective pressure on a specific myosin isoform in skeletal muscle may be different in each mammal. The plot of sequence vs mass for 2A (Fig 1) and 2X (Fig S1) do show more scatter than for β. It is remarkable therefore that our analysis points to a significant link between size and myosin sequence despite these factors.

In contrast the β-myosin is unusual in that there is only a single major slow isoform and therefore there is little scope to adjust the contraction parameters by mixing fiber types. The slow skeletal and cardiac muscle carrying β-myosin is fatigue resistant and also more economical with its energy usage, burning less ATP per unit of work output compared to the fast isoforms (Schiaffino and Reggiani 2011). Data on detailed biomechanics of contraction of muscles fibres expressing a single myosin isoform are not available for species outside standard lab animals. This makes a detailed assessment of selective pressure difficult. However, the heart and slow muscle fibres are relatively similar in function across mammals and the simple relationship between heart rate and size is well established. Only two myosin isoforms are expressed to any significant extent in the heart α and β, with α replacing β in the smallest mammals below ~ 1 kg. This may therefore allow the size-sequence divergence relationship to be more easily defined.

In most larger mammals (>1 kg) the α-myosin isoform is only expressed in the heart atrium where its role is to fill the ventricle and then β-myosin contracts the ventricle to deliver blood throughout the body. Thus α-myosin is a faster isoform and does not limit heart rate as the β-myosin does and this is likely to explain why sequence divergence of this isoform did not show a strong dependence on body mass (Table 1; Supplementary figure 1).

The median size of a species is linked to evolutionary success in quite complex ways. Cope’s rule (Stanley 1973; Hone and Benton 2005) for example states that in general species tend to evolve from the small to large, suggesting at the simplest level, a survival advantage of large over small individuals/species in competition for resources or mate selection. There are different factors that drive the need for larger animals to have slower contracting muscles. These include the mechanical constraints that require slower contracting muscles because of the larger mass to be moved and inertia. There may also be efficiency considerations as muscles are a significant source of heat generation and the balance between the need to keep warm and the need to lose heat alters between small and large animals. These two factors come together in the mammalian heart, where the basal metabolic rate dictates the rate at which the circulatory system delivers oxygen (plus other nutrients/waste products) and deals with heat dissipation between central and peripheral tissues and at the same time larger hearts beat more slowly as the volume of blood moved per contraction increases.

In summary, a negative correlation between species mass and heart rate is well established. We have presented evidence that the myosin motor domain of several of the adult muscle myosins have adapted to increasing species body mass to enable this reduction in contraction speed, particularly the β-myosin to enable reduced heart rate. Thus, providing a molecular explanation for the correlation of body mass and heart rate.

## Materials and Methods

Protein sequences were extracted from RefSeq (Pruitt et al. 2014) and Uniprot (UniProt Consortium 2014) as listed in Supplementary Table 1. Sequences were selected either as canonical, well annotated isoforms (e.g. human MyHC7 Uniprot:P12883), or by using BLAST (Altschul et al. 1990) to search for homologues of the human myosin proteins within the genomes of the other mammals. To ensure that each sequence corresponded to the requisite specific isoform, each sequence identified initially by database annotation or by BLAST, was further analysed using BLAST to compare it to UniProt to ensure that the sequence was complete and that it was most closely similar to the canonical standard for that isoform. Incomplete sequences were excluded from our analysis because sequence gaps could have a major effect on the sequence comparison for such closely related isoforms.

For each isoform, the protein sequences were divided into the Motor (1-800, β-myosin numbering) and Tail (842 – 1936) regions. For each region of each myosin, the sequences were aligned using Clustal Omega (Sievers et al. 2011); the resulting multiple sequence alignment was used to construct a percentage identity matrix between the species. Sequence identity was used rather than sequence similarity as we are considering small changes (>93% identity, >98% similarity) within the isoform and substitutions that would normally be classed as similar (e.g. aspartate to glutamate) may be relevant.

The masses of each species were extracted from a wide range of information sources and are listed in the Supplementary Information Tables 1. To compare sequence divergence against either evolutionary time or animal body mass, the relevant matrices were plotted against each other.

Both the protein sequence divergence and the dN/dS ratio were plotted against the species mass. In both cases, they had a robust regression fitted to reduce the weighting of the outliers. This is done by minimizing absolute difference rather than squared distance which should reduce the amount of under- and over-estimation in value difference caused by the square.

Residues associated with cardiomyopathies were extracted from UniProt and compared to the residues associated with body mass by considering exact matches and also whether residues associated with body mass were within +/− three residues of a residue associated with cardiomyopathy.

### Generation of phylogenetic trees

Two statistical methods were used to create phylogenetic trees, maximum likelihood (ML) and Bayesian inference methods, using programs Randomized Axelerated Maximum Likelihood (RAxML) (Stamatakis 2014) and Bayesian Evolutionary Analysis Sampling Trees (BEAST) and its corresponding user interface BEAUtI (Drummond et al. 2012), to increase reliability as species sequence identity is high. The RaxML trees were generated using the CIPRES Science Gateway (Miller et al.). TreeAnnotator (Drummond et al. 2012) was then used to generate a consensus tree for each set of 10000 trees produced by BEAST. *Monodelphis domestica* (opossum) was selected as an outgroup. The ML and Bayesian trees were compared using Compare2Trees (Nye, Liò, & Gilks, 2006), with black nodes on the trees indicating branches where the two trees disagree. Trees were drawn using FigTree (Rambaut. A - http://tree.bio.ed.ac.uk/software/figtree/). The patristic distances were calculated using the R “ape” package.

### Calculation of dN/dS variation

An analysis of the ratio of non-synonymous (dN) to synonymous (dS) variation for each sequence, when related to a potential evolutionary driving factor, can give an indication of positive selection. The dN/dS ratio was calculated pairwise to the *Mus musculus* (mouse) sequence by the yn00 program from the Phylogenetic Analysis by Maximum Likelihood (PAML) package (Yang 2007).

## Acknowledgements

MNW was supported by a Royal Society research grant. JM was supported by an EPSRC PhD studentship. MAG & JW were supported by funding from the European Union’s Horizon 2020 research and innovation programme under grant agreement No 777204. The authors would like to thank Martin Michaelis, Chris Toseland, and Carlo Reggiani for useful discussions and comments on the manuscript.

## Author Contributions

MNW, MAG and AJB devised the study. JEM, STJ, CH MPC and JW performed experiments. JEM, STJ, MNW, MAG, AJB analysed results. MR advised on use of statistics. All authors contributed to writing the manuscript.

## Data availability

The multiple sequence alignments used in this work are available from figshare – doi: 10.6084/m9.figshare.8362121

## References

Altschul SF, Gish W, Miller W, Myers EW, Lipman DJ. 1990. Basic local alignment search tool. J. Mol. Biol. 215:403–410.

Bloemink MJ, Geeves MA. 2011. Shaking the myosin family tree: biochemical kinetics defines four types of myosin motor. Semin. Cell Dev. Biol. 22:961–967.

Buvoli M, Hamady M, Leinwand LA, Knight R. 2008. Bioinformatics assessment of beta-myosin mutations reveals myosin’s high sensitivity to mutations. Trends Cardiovasc. Med. 18:141–149.

Drummond AJ, Suchard MA, Xie D, Rambaut A. 2012. Bayesian phylogenetics with BEAUti and the BEAST 1.7. Mol. Biol. Evol. 29:1969–1973.

Golomb E, Ma X, Jana SS, Preston YA, Kawamoto S, Shoham NG, Goldin E, Conti MA, Sellers JR, Adelstein RS. 2004. Identification and Characterization of Nonmuscle Myosin II-C, a New Member of the Myosin II Family. J. Biol. Chem. 279:2800–2808.

Goodson HV, Warrick HM, Spudich JA. 1999. Specialized conservation of surface loops of myosin: evidence that loops are involved in determining functional characteristics. J. Mol. Biol. 287:173–185.

Hill AV. 1950. The dimensions of animals and their muscular dynamics. Sci Prog 38:209–230.

Homburger JR, Green EM, Caleshu C, Sunitha MS, Taylor RE, Ruppel KM, Metpally RPR, Colan SD, Michels M, Day SM, et al. 2016. Multidimensional structure-function relationships in human ß-cardiac myosin from population-scale genetic variation. Proc. Natl. Acad. Sci. U.S.A. 113:6701–6706.

Hone DWE, Benton MJ. 2005. The evolution of large size: how does Cope’s Rule work? Trends Ecol. Evol. (Amst.) 20:4–6.

Johnson CA, McGreig JE, Vera CD, Mulvihill D, Ridout M, Leinwand LA, Wass MN, Geeves MA. 2019. Cardiac contraction velocity has evolved to match heart rate with body size through variation in β-cardiac myosin sequence. bioRxiv:680413.

Krendel M, Mooseker MS. 2005. Myosins: tails (and heads) of functional diversity. Physiology (Bethesda) 20:239–251.

Kurzawa-Goertz SE, Perreault-Micale CL, Trybus KM, Szent-Györgyi AG, Geeves MA. 1998. Loop I can modulate ADP affinity, ATPase activity, and motility of different scallop myosins. Transient kinetic analysis of S1 isoforms. Biochemistry 37:7517–7525.

Miller MA, Pfeiffer W, Schwartz T. Creating the CIPRES Science Gateway for inference of large phylogenetic trees. In: IEEE. pp. 1–8.

Moore JR, Leinwand L, Warshaw DM. 2012. Understanding cardiomyopathy phenotypes based on the functional impact of mutations in the myosin motor. Robbins J, Seidman C, Watkins H, editors. Circ. Res. 111:375–385.

Odronitz F, Kollmar M. 2007. Drawing the tree of eukaryotic life based on the analysis of 2,269 manually annotated myosins from 328 species. Genome Biol. 8:R196.

Pellegrino MA, Canepari M, Rossi R, D’Antona G, Reggiani C, Bottinelli R. 2003. Orthologous myosin isoforms and scaling of shortening velocity with body size in mouse, rat, rabbit and human muscles. The Journal of Physiology 546:677–689.

Pruitt KD, Brown GR, Hiatt SM, Thibaud-Nissen F, Astashyn A, Ermolaeva O, Farrell CM, Hart J, Landrum MJ, McGarvey KM, et al. 2014. RefSeq: an update on mammalian reference sequences. Nucleic Acids Res. 42:D756–D763.

Savage VM, Allen AP, Brown JH, Gillooly JF, Herman AB, Woodruff WH, West GB. 2007. Scaling of number, size, and metabolic rate of cells with body size in mammals. Proc. Natl. Acad. Sci. U.S.A. 104:4718–4723.

Schiaffino S, Reggiani C. 2011. Fiber types in mammalian skeletal muscles. Physiol. Rev. 91:1447–1531.

Sievers F, Wilm A, Dineen D, Gibson TJ, Karplus K, Li W, Lopez R, McWilliam H, Remmert M, Söding J, et al. 2011. Fast, scalable generation of high-quality protein multiple sequence alignments using Clustal Omega. Mol. Syst. Biol. 7:539.

Spudich JA. 1994. How molecular motors work. Nature 372:515–518.

Stamatakis A. 2014. RAxML version 8: a tool for phylogenetic analysis and post-analysis of large phylogenies. Bioinformatics 30:1312–1313.

Stanley SM. 1973. An Explanation for Cope’s Rule. Evolution 27:1.

Sweeney HL, Rosenfeld SS, Brown F, Faust L, Smith J, Xing J, Stein LA, Sellers JR. 1998. Kinetic tuning of myosin via a flexible loop adjacent to the nucleotide binding pocket. J. Biol. Chem. 273:6262–6270.

UniProt Consortium. 2014. Activities at the Universal Protein Resource (UniProt). Nucleic Acids Res. 42:D191–D198.

Walsh R, Rutland C, Thomas R, Loughna S. 2010. Cardiomyopathy: a systematic review of disease-causing mutations in myosin heavy chain 7 and their phenotypic manifestations. Cardiology 115:49–60.

Yang Z. 2007. PAML 4: phylogenetic analysis by maximum likelihood. Mol. Biol. Evol. 24:1586–1591.

