## Supplementary Figures for "Adaptation of mammalian myosin II sequences to body mass"

**Supplementary Figure 1. Mass vs sequence identity plots.** The motor and tail domains have been analysed separately. The grey squares are Euarchontoglires, the black triangles are Laurasiatheria and the open circles are the Afrotheria and Metatheria groups. Each plot has been fitted with a robust linear regression. Sequence identity is pairwise to the mouse. The $R^2$ value and slope gradient are shown on each plot.

**A – Tail domains for EMB, β, NMA, 2B, NMB and 2A**

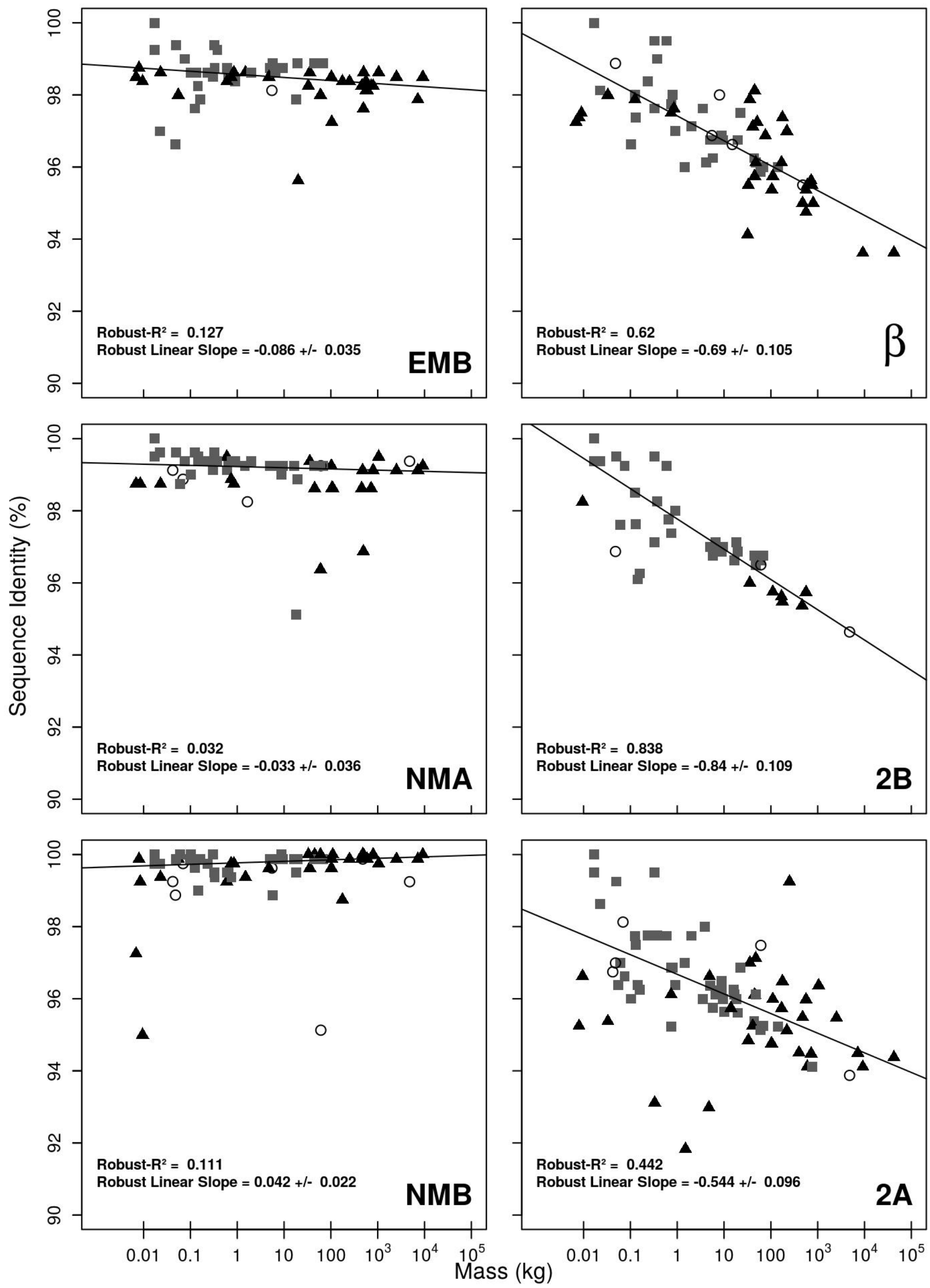

**B – Motor and Tail domains for 2X, EXOC and SM**

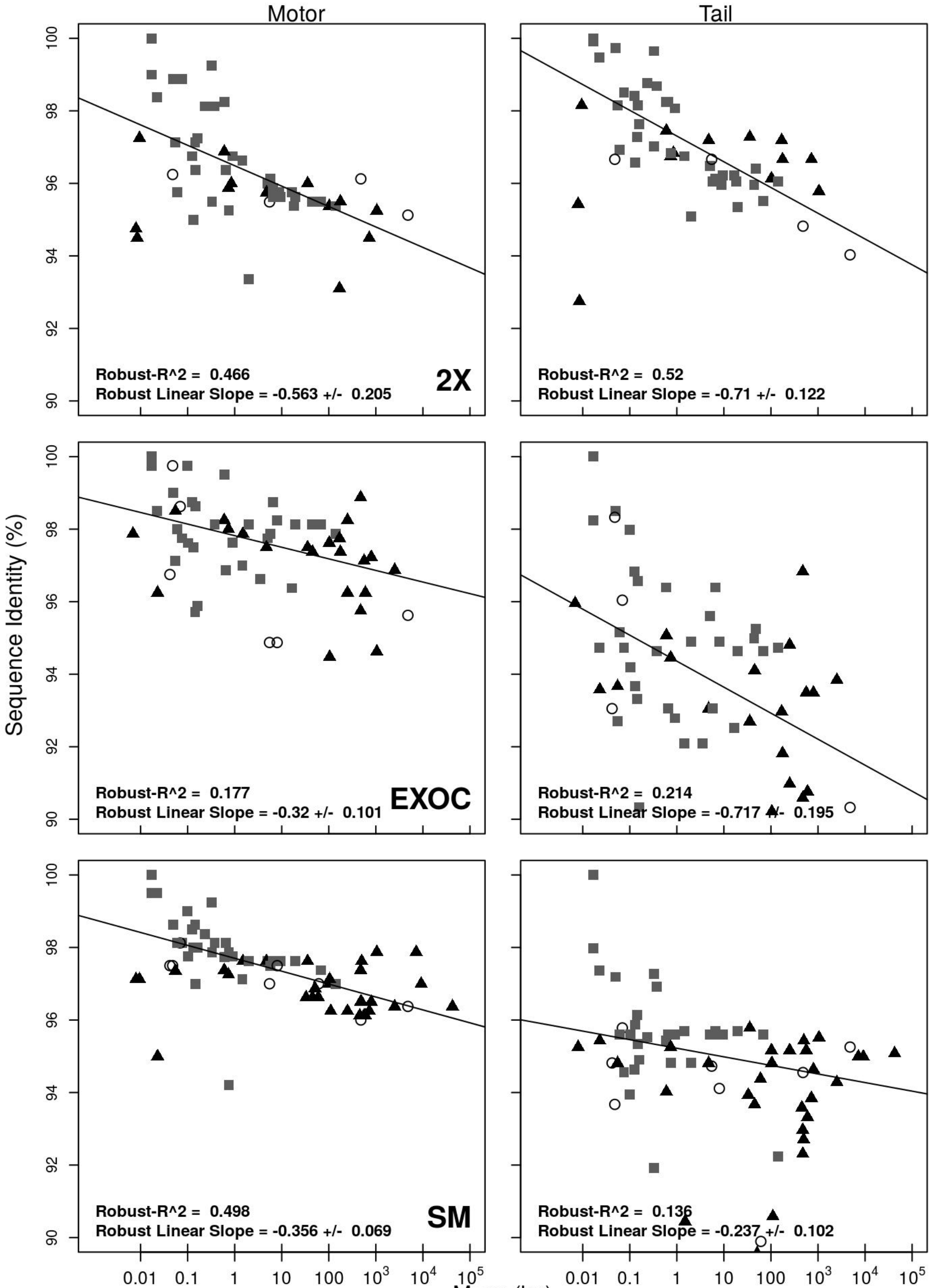

**C – Motor and Tail domains for PERI, SlowT, and α**

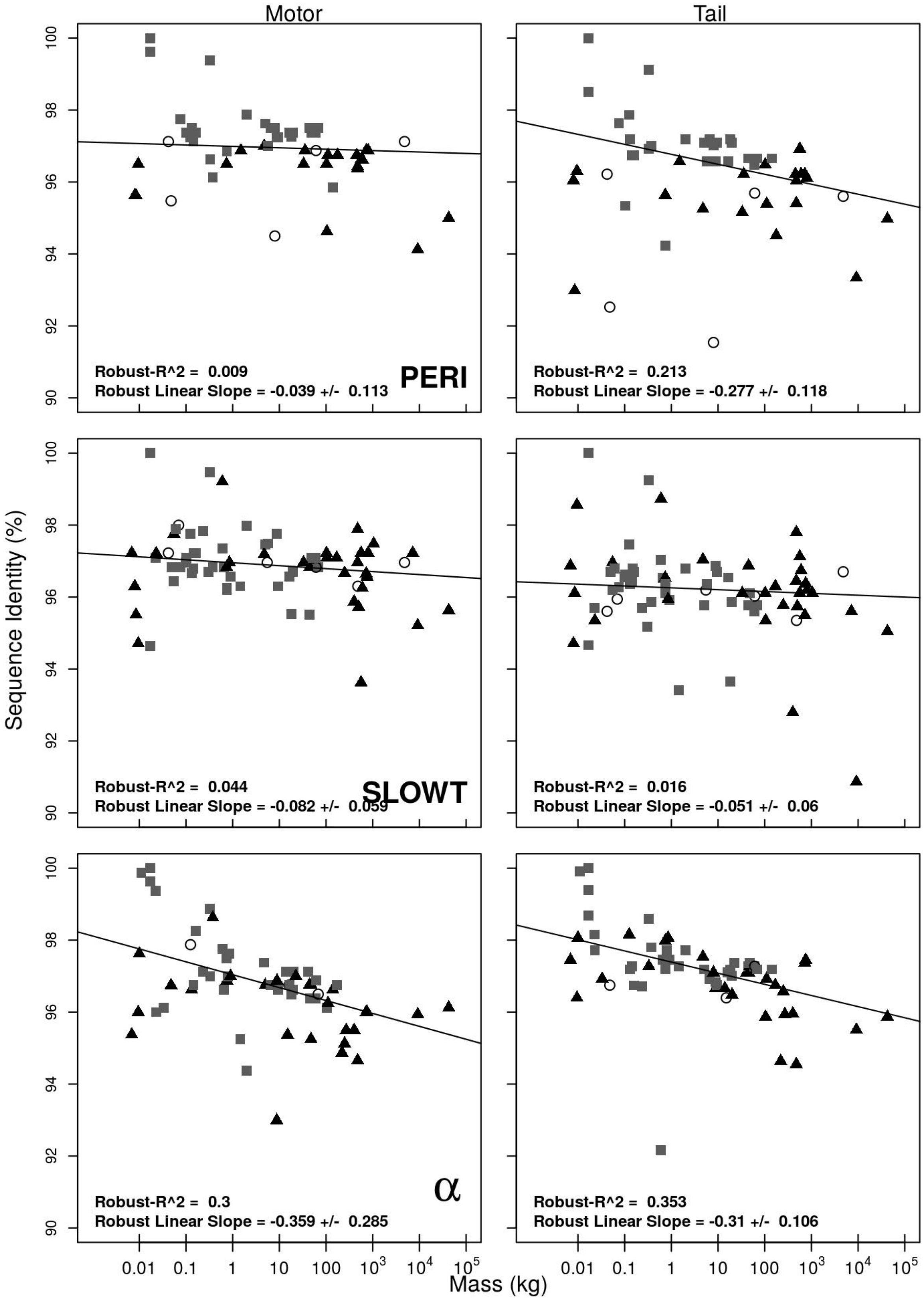

**Supplementary Figure 2. Phylogenetic Tree for the coding DNA sequences of EMB and NMA motor domain.**

Blue lines are Euarchontoglires, red lines are Laurasiatheria and black lines are anything else. The black circles at the nodes indicate where different grouping was seen in trees generated by Maximum likelihood and Bayesian methods.

**A – EMB**

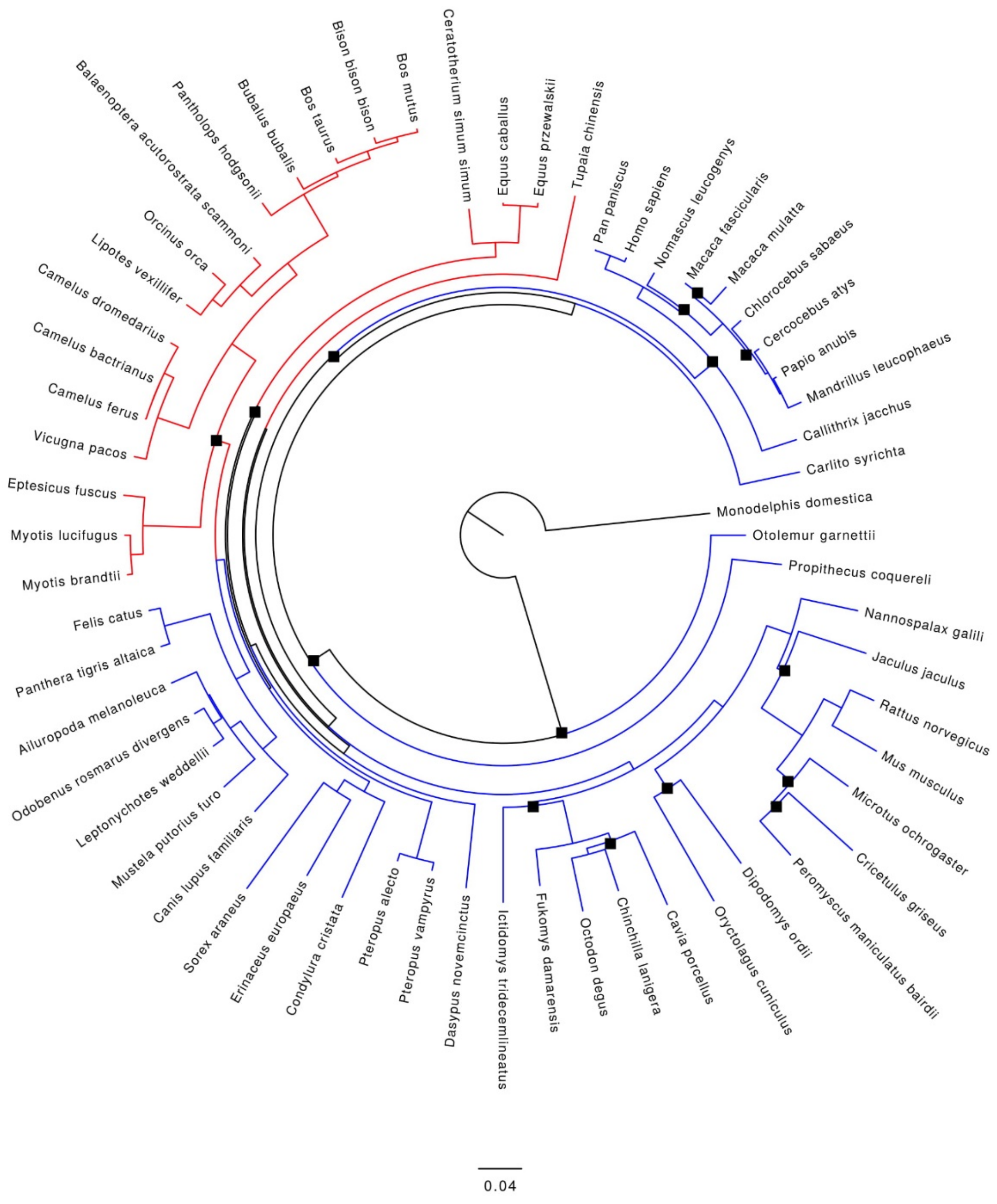

**B – NMA**

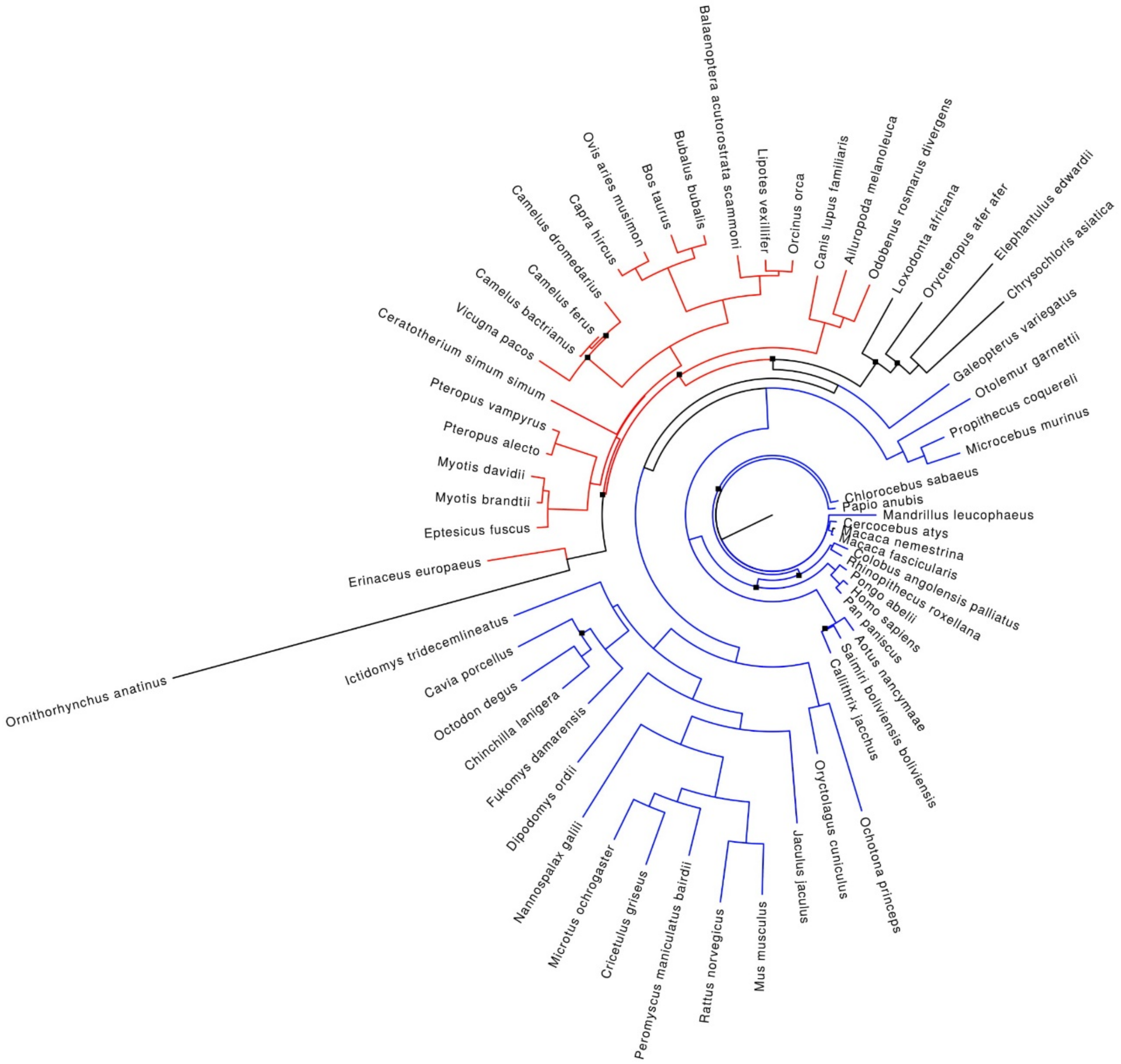

Balaenoptera acutorostrata scammoni
Lipotes vexillifer
Orcinus orca
Canis lupus familiaris
Ailuropoda melanoleuca
Odobenus rosmarus divergens
Loxodonta africana
Orycteropus afer afer
Elephantulus edwardii
Chrysochloris asiatica
Galeopterus variegatus
Otolemur garnettii
Propithecus coquereli
Microcebus murinus
Chlorocebus sabaeus
Papio anubis
Mandrillus leucophaeus
Cercocebus atys
Macaca nemestrina
Macaca fascicularis
Colobus angolensis palliatus
Rhinopithecus roxellana
Pongo abelii
Homo sapiens
Pan paniscus
Aotus nancymaae
Saimiri boliviensis boliviensis
Callithrix jacchus
Ochotona princeps
Oryctolagus cuniculus
Jaculus jaculus
Mus musculus
Rattus norvegicus
Peromyscus maniculatus bairdii
Cricetulus griseus
Microtus ochrogaster
Nannospalax galili
Dipodomys ordii
Fukomys damarensis
Chinchilla lanigera
Octodon degus
Cavia porcellus
Ictidomys tridecemlineatus
Ornithorhynchus anatinus
Erinaceus europaeus
Eptesicus fuscus
Myotis brandtii
Myotis davidii
Pteropus alecto
Pteropus vampyrus
Ceratotherium simum simum
Vicugna pacos
Camelus bactrianus
Camelus ferus
Camelus dromedarius
Capra hircus
Ovis aries musimon
Bos taurus
Bubalus bubalis

**Supplementary Figure 3. Patristic distances as a function of mass.** The patristic distance for each species in the ML trees in relation to the mouse sequence for EMB, β, and NMA as a function of species mass. The grey squares are Euarchontoglires, the black triangles are Laurasiatheria and the open circles are the Afrotheria and Metatheria groups.

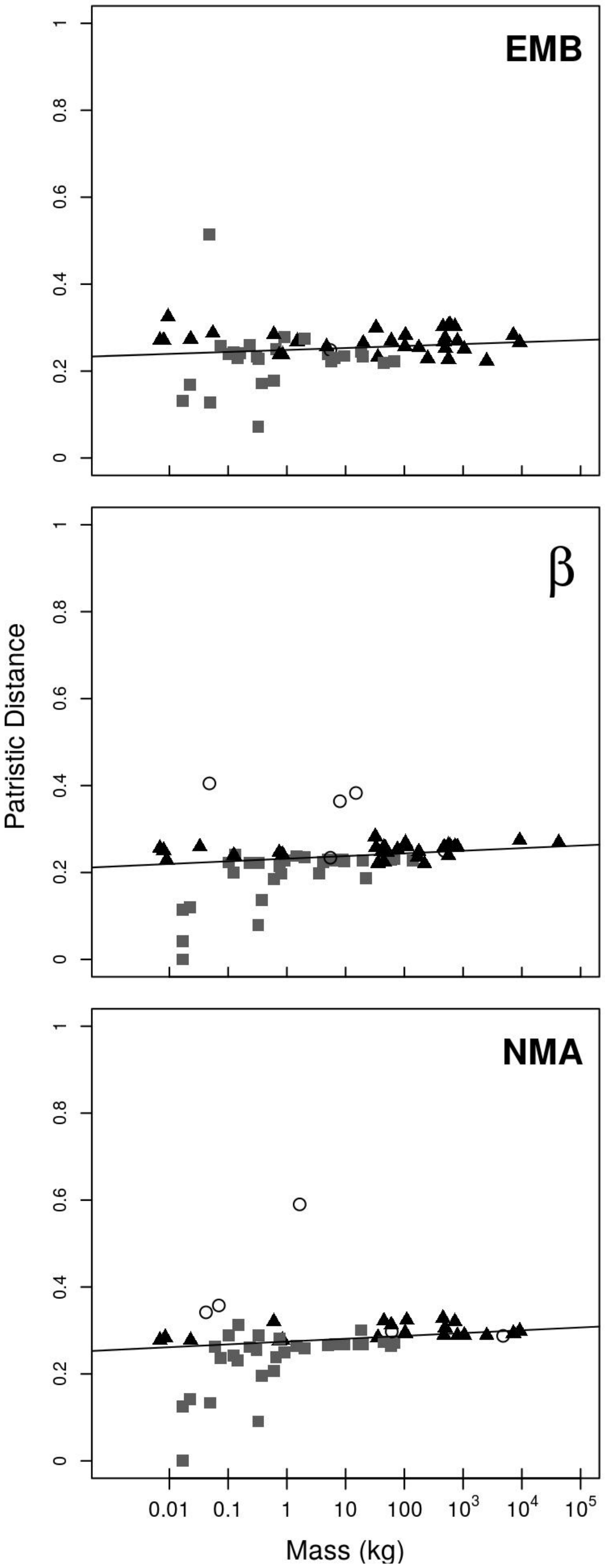

**Supplementary Figure 4. The indication of positive selection on species mass for three myosin heavy chains.** Non-synonymous (dN) and synonymous (dS) data for each species compared to the mouse sequence for EMB, β, and NMA as a function of species mass. Details of the species mass and sequences used are given in Supplementary Table 1. The grey squares are Euarchontoglires, the black triangles are Laurasiatheria and the open circles are the Afrotheria and Metatheria groups.

**A – dN for EMB, β, and NMA**

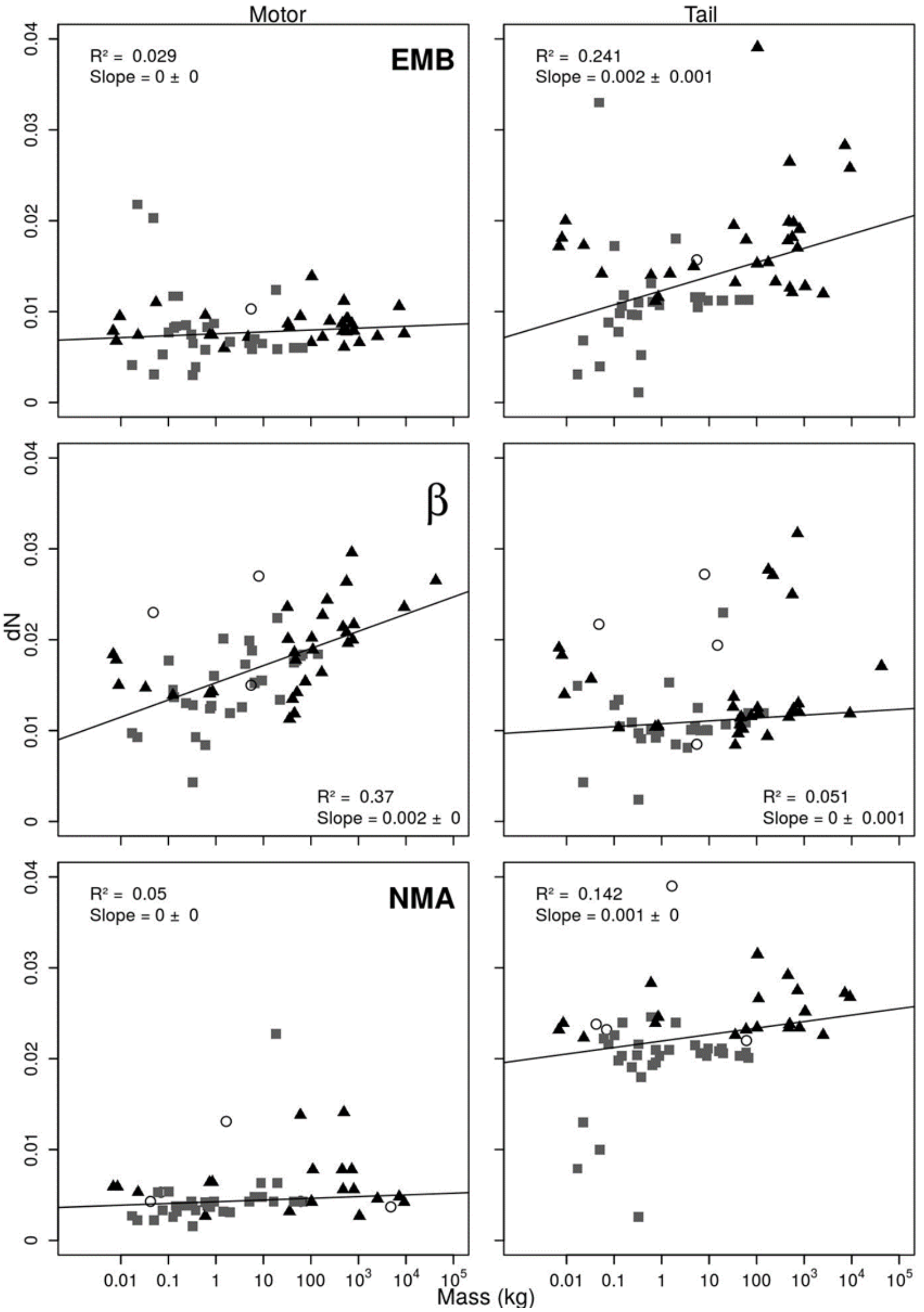

**B – dS and dN/dS for EMB, β, and NMA Tail regions**

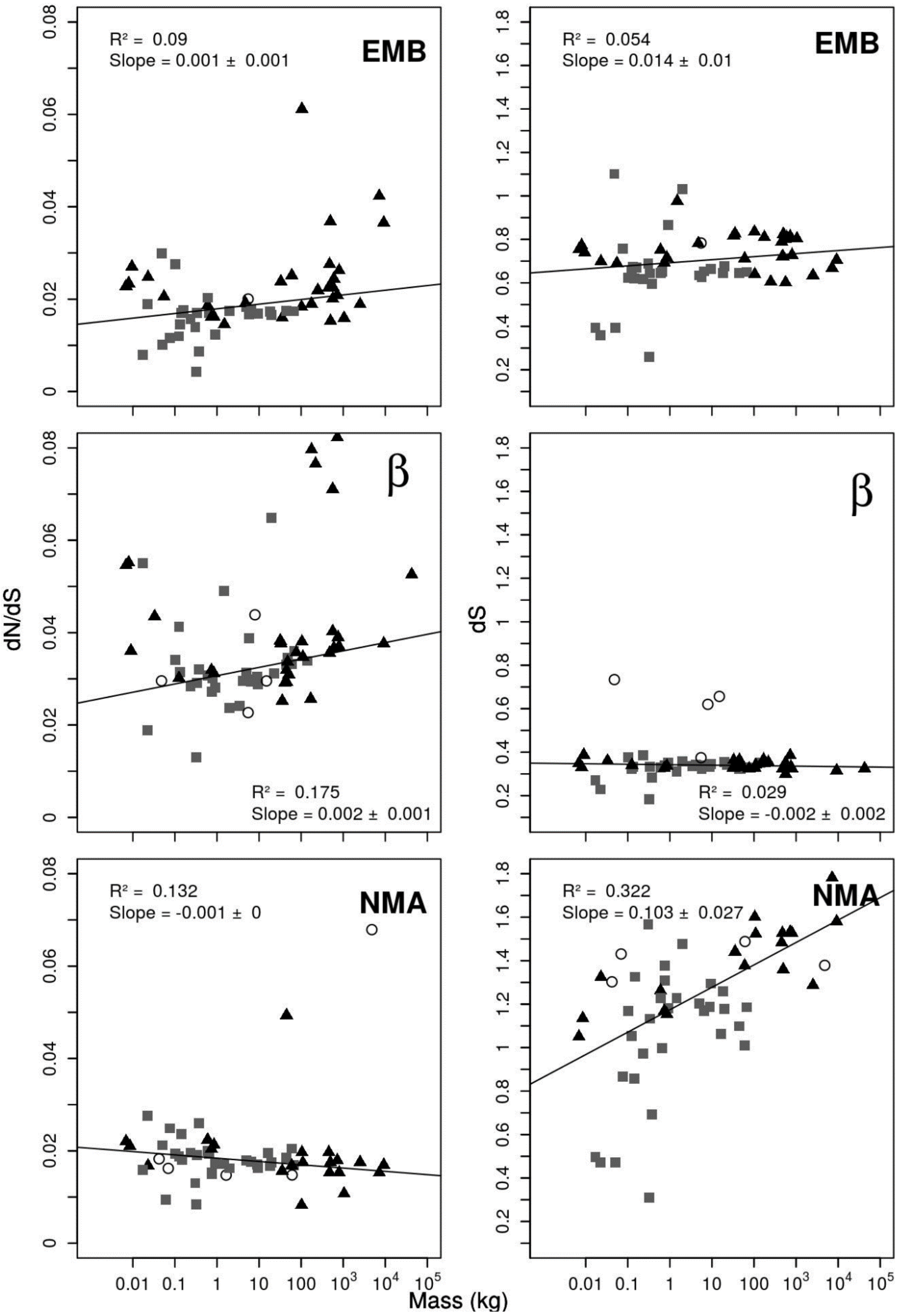

**Supplementary Figure 5. The indication of positive selection on species mass for adult sarcomeric myosin II isoforms.** Non-synonymous (dN), synonymous (dS) and dN/dS ratio data for each species compared to the mouse sequence for the adult sarcomeric Myosin II isoforms as a function of species mass. Details of the species mass and sequences used are given in Supplementary Table 1. The grey squares are Euarchontoglires, the black triangles are Laurasiatheria and the open circles are the Afrotheria and Metatheria groups.

**A – dN for 2B, 2A, 2X, and α**

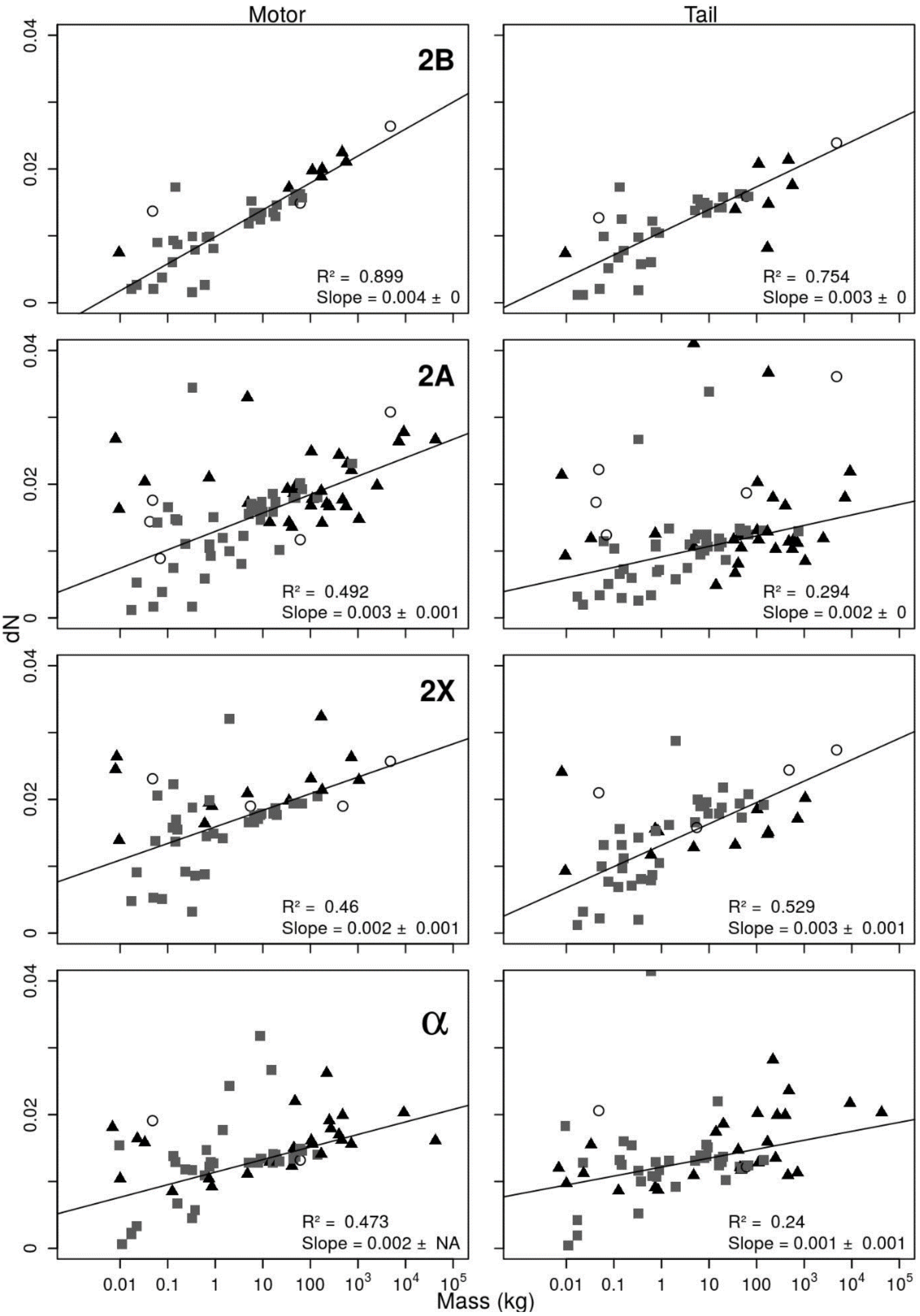


**B – dS for 2B, 2A, 2X, and α**

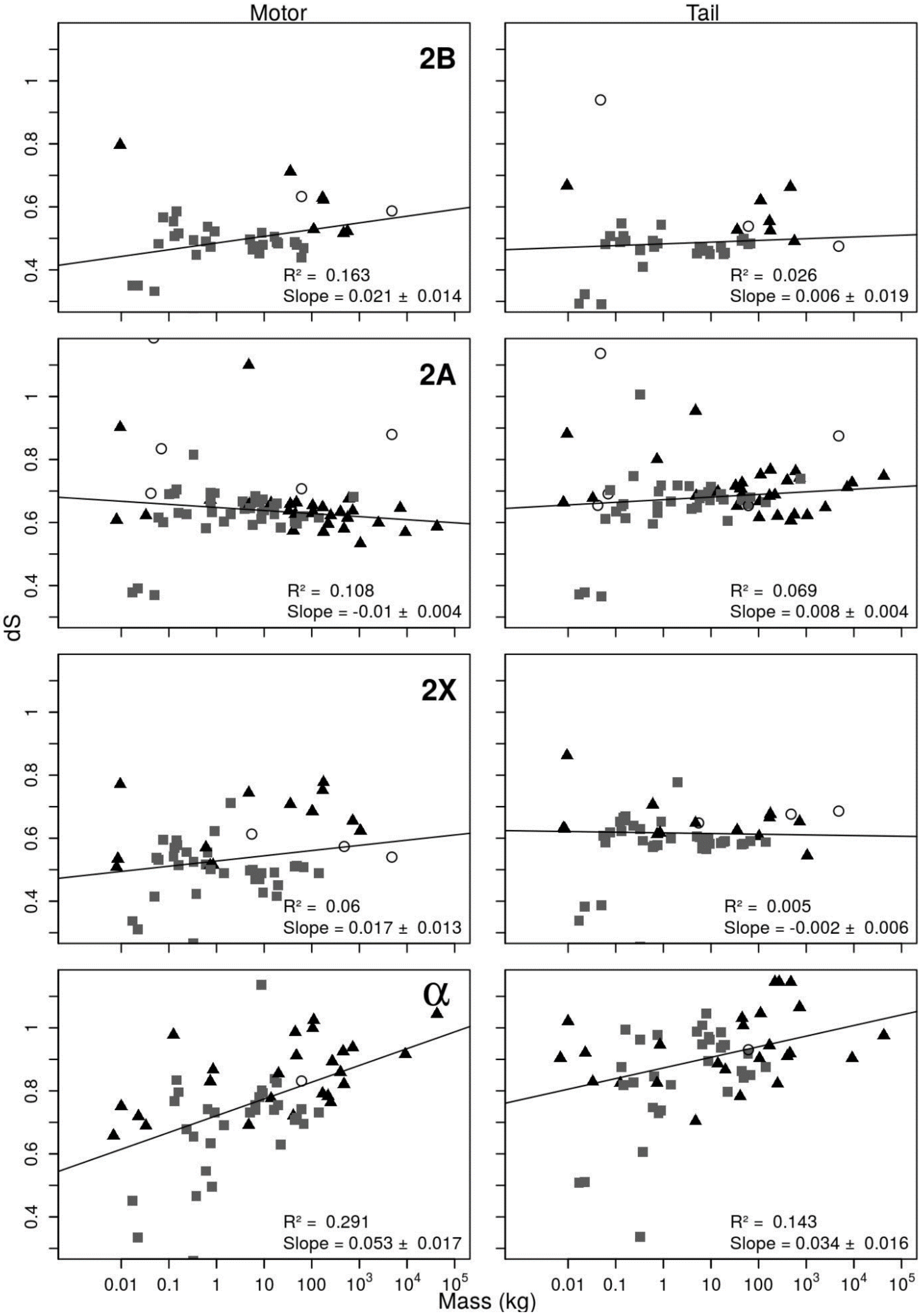

Motor
Tail
2B
R² = 0.163
Slope = 0.021 ± 0.014
R² = 0.026
Slope = 0.006 ± 0.019
2A
R² = 0.108
Slope = -0.01 ± 0.004
R² = 0.069
Slope = 0.008 ± 0.004
2X
R² = 0.06
Slope = 0.017 ± 0.013
R² = 0.005
Slope = -0.002 ± 0.006
α
R² = 0.291
Slope = 0.053 ± 0.017
R² = 0.143
Slope = 0.034 ± 0.016
dS
Mass (kg)

**C – dN/dS for 2B, 2A, 2X, and α**

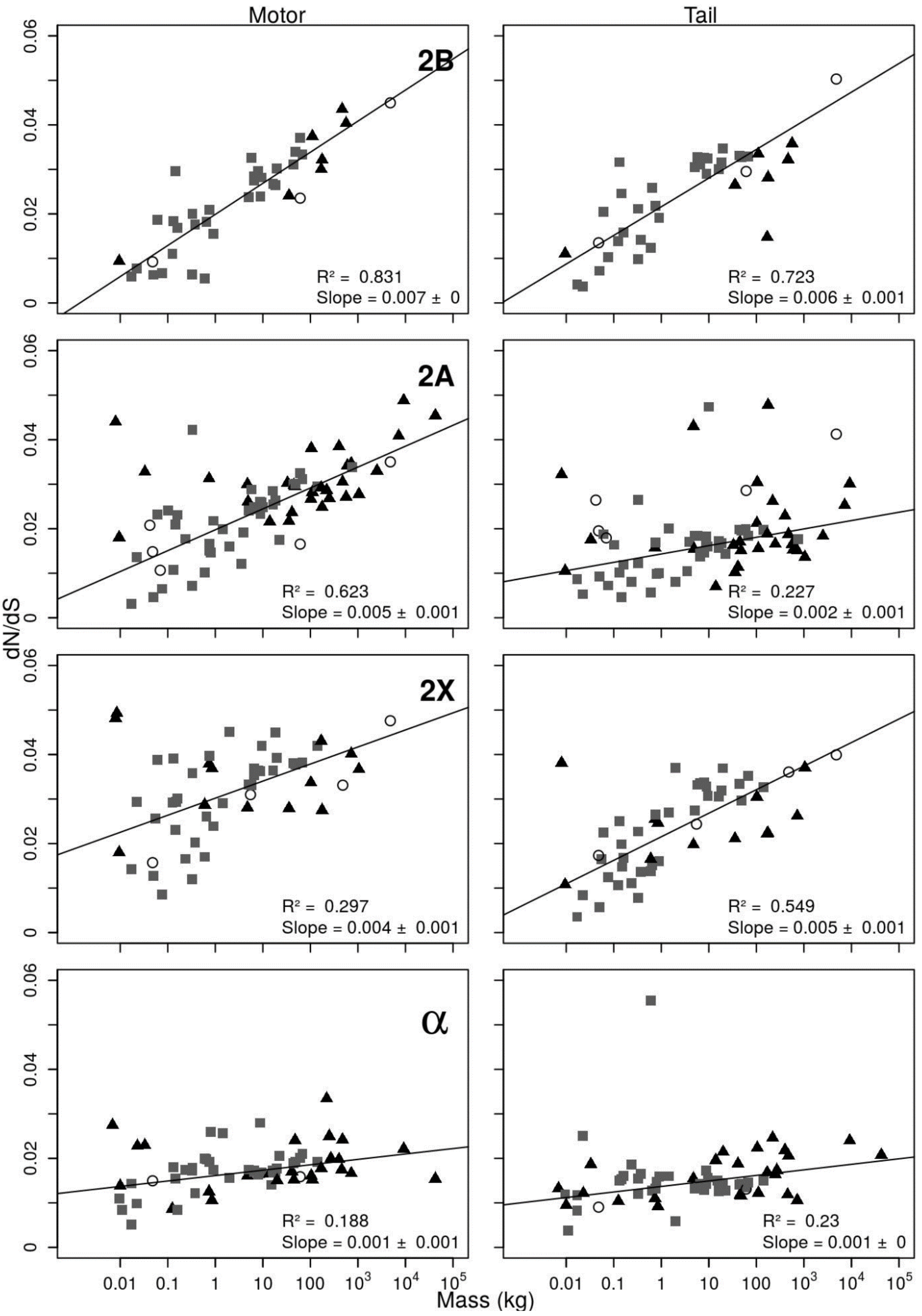

**Supplementary Figure 6. Plot conservation for EMB, 2A, 2B, and NMA motor and tail regions.**

Blue lines show the frequency of missense mutations at their corresponding residue position for human EMB and NMA present in the Genome Aggregation Database (gnomAD). Red lines show the number of times the consensus amino acid occurs at each residue position in the sequence. Black lines show the number of different amino acids occurring at that position in the sequence (minimum 1). The heads (A, C) and tails (B, D) are shown in different plots to allow an expanded scale for the head domain.

**A – EMB Motor**

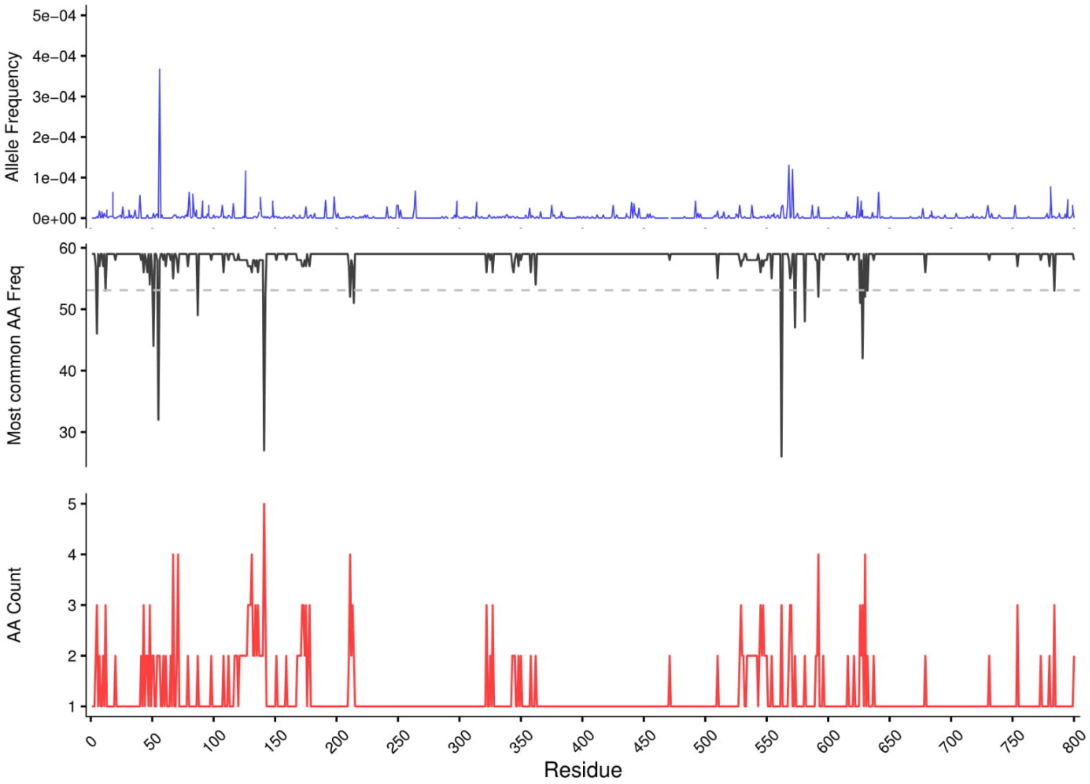


**B – EMB Tail**

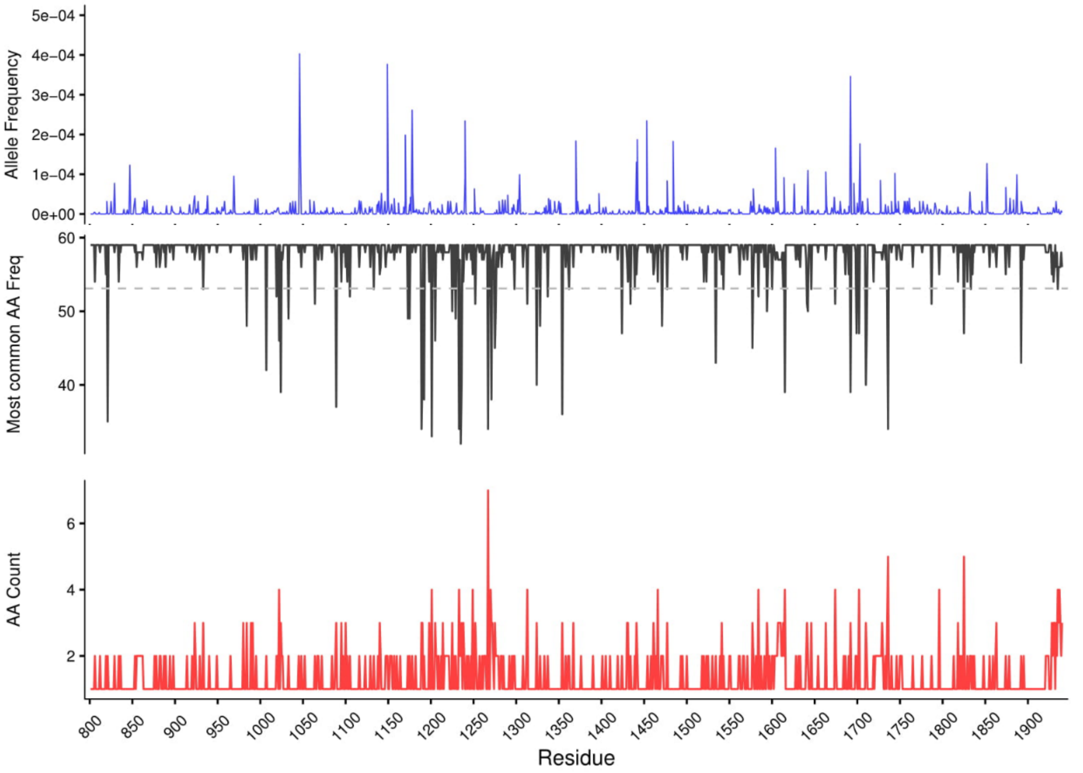

C – NMA Motor

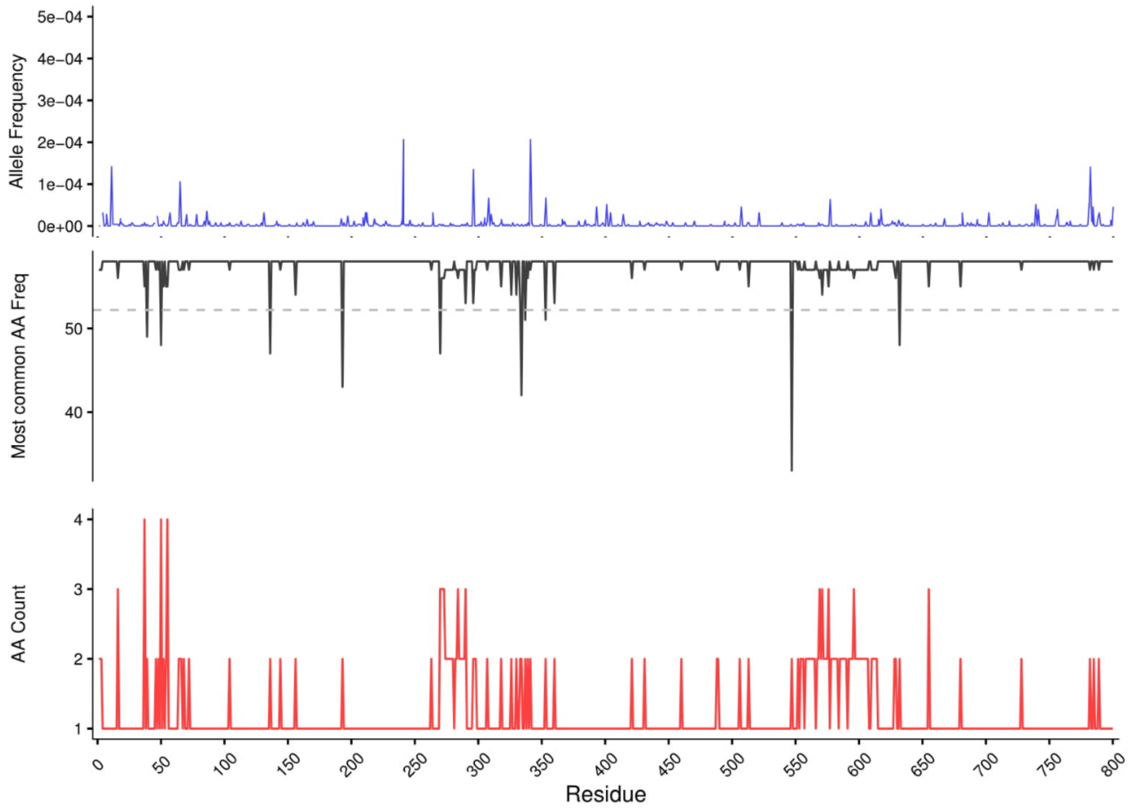


D – NMA Tail

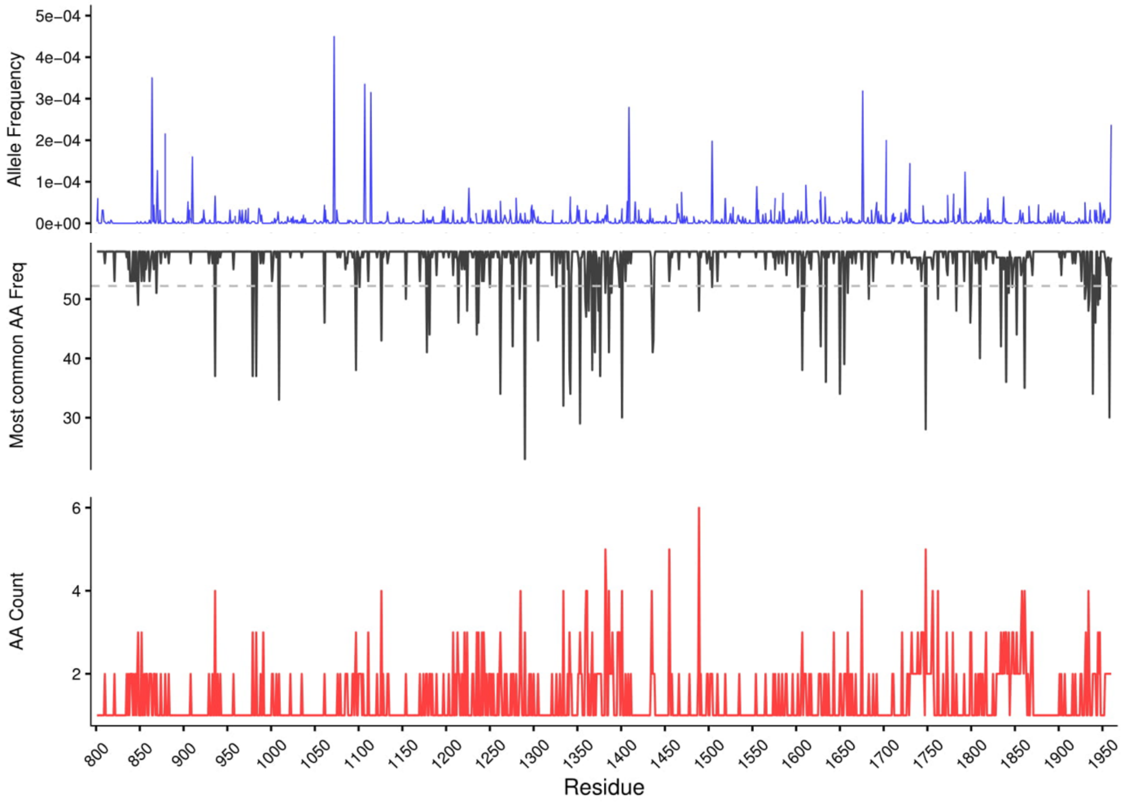

### E – 2B Motor and Tail

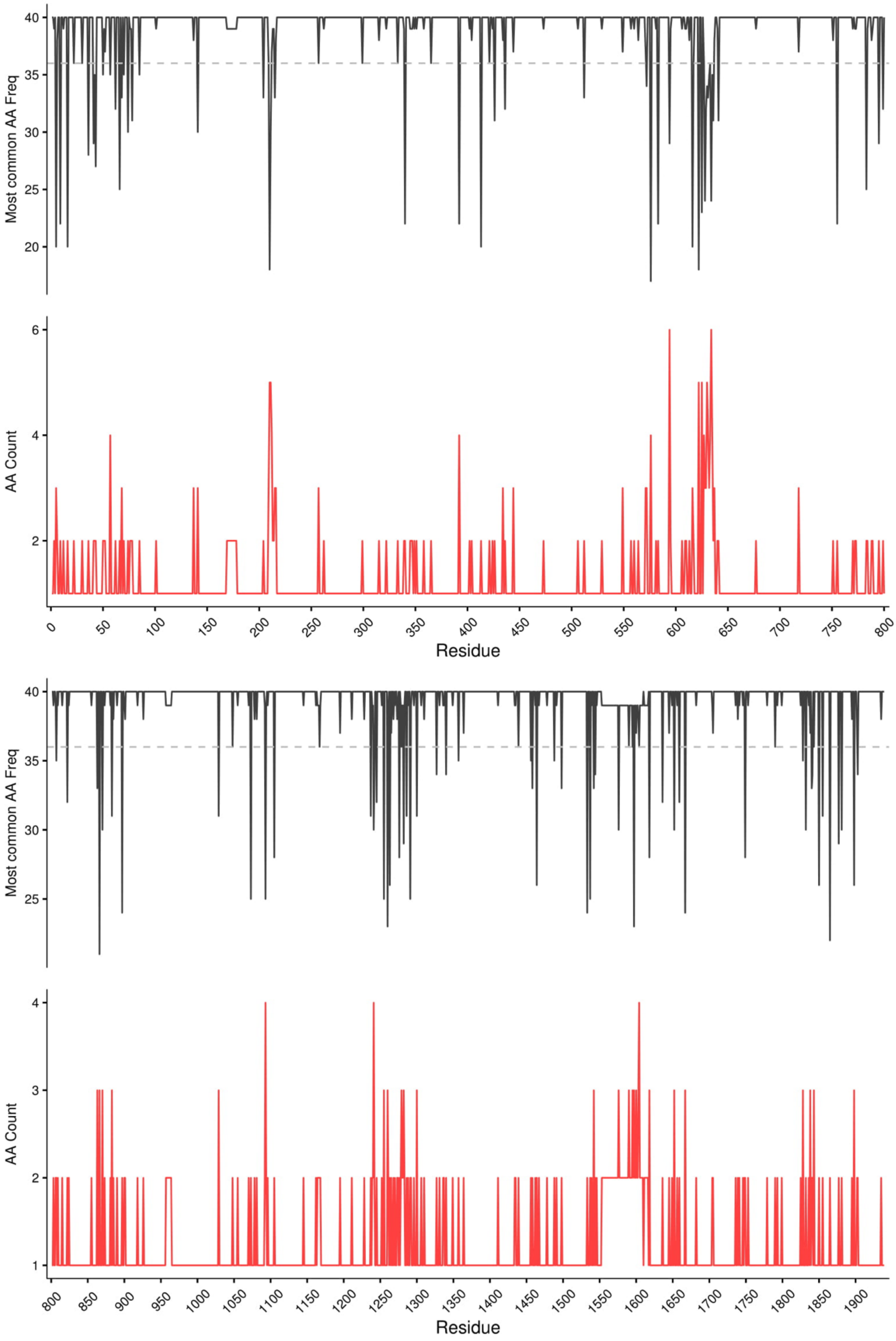

### F – 2A Motor and Tail

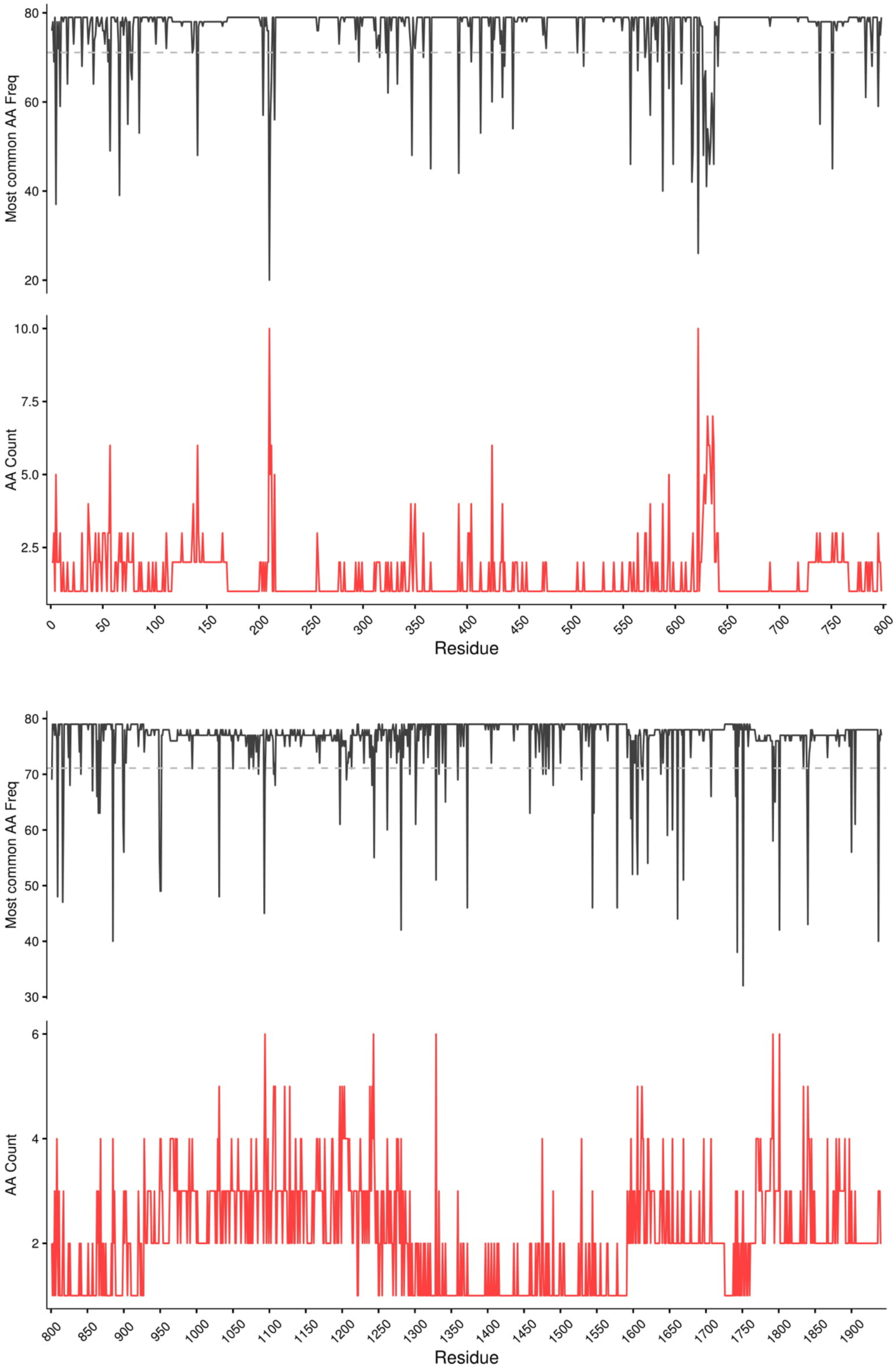
